# Surprising connections between DNA binding and function for the near-complete set of yeast transcription factors

**DOI:** 10.1101/2023.07.25.550593

**Authors:** Lakshmi Mahendrawada, Linda Warfield, Rafal Donczew, Steven Hahn

## Abstract

DNA sequence-specific transcription factors (TFs) modulate transcription and chromatin architecture, acting from regulatory sites in enhancers and promoters of eukaryotic genes. How TFs locate their DNA targets and how multiple TFs cooperate to regulate individual genes is still unclear. Most yeast TFs are thought to regulate transcription via binding to upstream activating sequences, situated within a few hundred base pairs upstream of the regulated gene. While this model has been validated for individual TFs and specific genes, it has not been tested in a systematic way with the large set of yeast TFs. Here, we have integrated information on the binding and expression targets for the near-complete set of yeast TFs. While we found many instances of functional TF binding sites in upstream regulatory regions, we found many more instances that do not fit this model. In many cases, rapid TF depletion affects gene expression where there is no detectable binding of that TF to the upstream region of the affected gene. In addition, for most TFs, only a small fraction of bound TFs regulates the nearby gene, showing that TF binding does not automatically correspond to regulation of the linked gene. Finally, we found that only a small percentage of TFs are exclusively strong activators or repressors with most TFs having dual function. Overall, our comprehensive mapping of TF binding and regulatory targets have both confirmed known TF relationships and revealed surprising properties of TF function.

## Introduction

The subset of DNA sequence-specific transcription factors (TFs) expressed in a eukaryotic cell determines its fundamental properties including gene expression patterns, cell identity, and the responses to both intrinsic signals and environmental changes^1–6^. TFs primarily act by directing chromatin modifications, including higher order chromatin structure, and/or modulating transcription initiation or elongation via direct interactions with cofactors or the basal transcription machinery^3, 7^. TF-DNA interactions are highly dynamic^8^ and the mechanisms by which TFs locate their specific DNA targets within the genome remains an open question. Many eukaryotic TFs recognize short DNA binding motifs; however, many motifs seem to lack sufficient information to uniquely specify functional sites in large genomes^9, 10^. TF-DNA interactions mediated by sequence-dependent DNA structure is another well-supported mechanism^3, 9, 11^. Other alternative pathways for TF binding include recruitment via protein-protein interactions with DNA-bound TFs, recruitment mediated by disordered regions of the TFs^12, 13^ or by cooperative binding of TFs that generate unique DNA binding specificity^14, 15^.

While some TFs activate transcription^16–18^, recent large-scale screens suggest that only a small fraction of mammalian TFs possess strong activation function in the absence of other TFs^16, 19^, suggesting that most TFs must act in combination to regulate transcription. Several models have been proposed to explain how functional gene regulatory elements such as enhancers arise from combining multiple TF binding sites. In the interferon-beta enhanceosome, activity is strictly dependent on the order and orientation of the TF binding sites, resulting in highly cooperative binding^20^, however, this mechanism seems relatively rare. In contrast, the billboard model^15, 23^ proposes that TFs work together in a combinatorial manner without strict binding site grammar ^11, 21^. Another model suggests that certain TFs have preferences to function together, and that the order and orientation of binding sites moderately affects enhancer function^15, 22^. According to this latter model, the combined activity of the element is moderately higher than the sum of the individual TF activities. Recent large-scale screens support many but not all genes being regulated by the latter two models. For example, a large-scale screen for UAS activity using random sequences suggested that many individual TF motifs have only weak effects on gene expression^23^. This implied that, for many genes, enhancer activity is generated by combining modest activities of individual TFs. However, many studies often draw conclusions about TF function and specificity based on TF motifs present or absent in regulatory regions rather than directly assessing TF binding. Since at least some in vivo binding appears to be motif-independent, this approach may bias conclusions about the function and cooperativity of individual TFs.

Promoters of *S. cerevisiae* (yeast) protein-coding genes have been classified based on cofactor dependence for factors such as TFIID, Bdf1/2, SAGA, and the activator-binding domain of Mediator termed Tail^24–26^. Transcription from the majority of yeast genes is dependent on TFIID (TFIID-dependent genes) and many of these genes are constitutively expressed, while transcription from a smaller subset of mostly regulated genes, designated as coactivator redundant (CR), can use either TFIID or SAGA. Recent findings suggest that cofactor specificity of CR genes is determined by the identity of both UAS and core promoter sequences, while cofactor specificity of TFIID-dependent genes is defined by the core promoter^27^. How TFs contribute to cofactor specificity and whether certain TFs are specific for particular gene classes are important open questions. For example, it’s been estimated that about 60% of yeast TFs contain an acidic activation domain of the type that binds MED Tail^17, 18^, yet transcription from only ∼ 6% of genes is sensitive to rapid Tail depletion^25^.

Yeast is an excellent model system for studying TF specificity, cooperativity, and function. Yeast shares many transcription mechanisms with higher eukaryotes, albeit lacking known long-range enhancers^2^. Yeast have only ∼ 150 total TFs, about ten times fewer than humans, making it suitable for comprehensive large-scale binding and functional comparisons. A wealth of genetic and biochemical data is available on many TF-TF interactions and on TFs that co-regulate sets of genes^28–31^. However, recent formaldehyde-based crosslinking analysis of 78 yeast TFs surprisingly suggested that only ∼20% of yeast genes have evolved to be regulated by TFs and that the other gene sets either had no detectable TFs (46%) or only chromatin organizing/insulator factors^30^.

Here we took an alternative approach to probe the genome-wide roles of yeast TFs, mapping both the DNA locations and expression targets for 126 TFs. First, we observed that most Pol-II promoters are occupied by at least one TF. Second, while we found that nearly all genes are regulated by one or more TFs, our analysis revealed surprisingly limited overlap between TF binding and gene regulation. Third, rapid depletion of many TFs revealed numerous regulatory targets distant from detectable TF binding sites, suggesting long-range and/or chromatin-mediated regulatory mechanisms. Our study provides a comprehensive survey of TF functions, offering important insights into interactions between the set of TFs expressed in a single cell type and how they contribute to the complex mechanisms of gene regulation.

## Results

### Genome-wide binding patterns of yeast transcription factors

To understand the roles and coordination of transcription factor (TF) function at a systems-wide level, we first mapped the binding locations for nearly all yeast sequence-specific TFs. This set includes transcription activators, repressors, and putative transcription factors. For comparison, we also mapped the binding for some chromatin remodelers, transcription cofactors, and preinitiation complex (PIC)- associating proteins (**Table S1**). Strains were created for each of 212 proteins individually fused to MNase and their positions on genomic DNA assayed using ChEC-seq, where DNA cleavage directed by each TF-MNase fusion maps its chromosomal locations ^26, 32, 33^. Unless otherwise stated, TF-DNA binding was assayed in cells grown in synthetic complete glucose media and all assays were performed in triplicate. Our stringent peak calling criteria requires both a 10-fold higher TF-MNase cleavage compared with a free MNase control and a 6-fold signal above the 10 kb local background (Methods).

From this analysis, 178 factors exhibited consistent and statistically significant binding to at least one promoter/regulatory region of protein coding genes (defined as −400 bps upstream and 200 bps downstream from TSS), hereafter termed “promoters” for simplicity (**Table S1, Fig 1A & S1A**). We found that 5467 out of 5891 assayed promoters detectably interact with at least one TF and a typical promoter interacts with a median of 15 TFs but ranges from 1-137 TFs/promoter (**Table S2**). The frequency distribution of TF binding sites showed that most TFs target < 1000 promoters with a median of 624 promoters targeted per TF but ranges from 1-3355 promoters/TF (**Fig S1B**). Prior studies demonstrated that many highly expressed inducible promoters are enriched in the CR gene class, while constitutive promoters are mostly in the TFIID class ^26, 34^ and we investigated whether any TFs exhibit class-specific binding patterns. Some TFs (16) display significant enrichment at TFIID promoters beyond random expectation (p-value < 0.05; hypergeometric test) (e.g., YLL054C in **Fig S1C**) while some TFs (18) have no preference for either class (e.g., Fhl1 in **Fig S1C**). Strikingly, CR promoters interact with a much greater number of TFs (**Fig S1D**), and many of these TFs (144; p-value < 0.05; hypergeometric test) showed selective enrichment at CR promoters (e.g., Mga1 in **Fig S1C**). These findings support the notion that genes displaying high transcriptional variance across different growth or stress conditions have a more complex network of TF control compared with constitutive genes.

**Figure 1.**
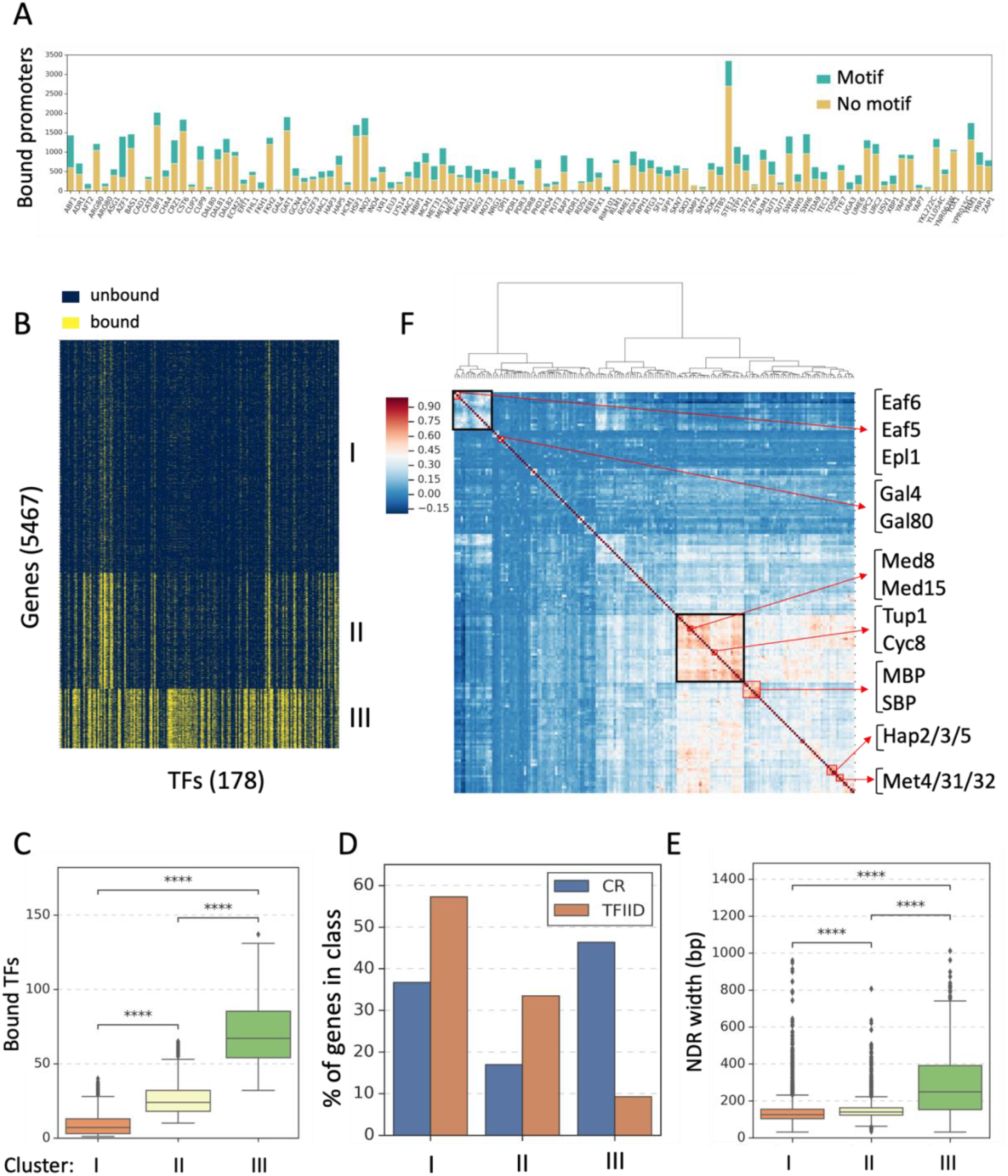
Genome wide mapping of yeast transcription factor binding. **(A)** For 107 yeast transcription factors (TFs) with DNA sequence motifs listed in the Jaspar database^35^, the stacked bar plot depicts experimentally observed TF binding sites in regulatory regions of protein coding genes with and without the known DNA sequence motif (green and yellow bars respectively). FIMO^36^ was used to scan binding sites for the presence of motifs (Methods) with a threshold of p < 0.00025. DNA sequences of all promoters (−400 to +200 bps from TSS) were used as the background model. **(B)** The heatmap represents binary binding events of 178 TFs to 5467 promoters clustered by unsupervised K-means. Yellow bars represent binding and dark blue no binding. **(C)** Box plot shows the number of TFs detected at gene regulatory regions in each cluster from panel 1B: Cluster-I (1-40 TFs); Cluster-II (10-65 TFs); Cluster-III (32-137 TFs). Results of Welch’s t-test are shown in 1C-1E. Significance for this and for all subsequent figures is defined as -ns: > 0.05, *: 0.05-0.01, **: 0.01-0.001, ***: 0.001-0.0001, ****: p <= 0.0001. **(D)** The distribution of our TF binding clusters (Fig 1B) among TFIID and CR genes^26^ is shown. **(E)** Box plot shows the NDR width of promoters in each cluster. NDR width has been reported for 5237 out of 5467 analyzed promoters^37^. **(F)** Correlation between TFs based on binding events. Cluster map displays the hierarchical clustering of TF-TF correlations. Top left cluster highlighted in black contains TFs that are enriched at Cluster-II genes; middle cluster highlighted in black contains TFs enriched in Cluster-III genes. Examples of previously established TF interactions are highlighted in red. Correlation values range from −0.15 to 0.9.

To assess DNA sequence specificity of TF binding sites, we analyzed TF ChEC-seq peak locations for the presence of consensus DNA binding motifs using FIMO^36^, with promoter sequences (400 bps upstream and 200 bps downstream of all 5891 TSS) of all genes used as a background model. Relatively few TFs (23 TFs, e.g., Abf1, Reb1) contained a consensus motif at >50% of their binding sites while, for many other TFs, only a fraction of binding sites (median − 30%; range − 5 to 48%) contained a consensus motif (**Fig 1A**). While somewhat unexpected, this finding agrees with analysis of published data where formaldehyde-based crosslinking was used to map a smaller set of yeast sequence specific TFs (**Fig S2A**)^30^ and with the findings that, for some TFs, disordered regions outside of the DNA binding domain can dictate targeting to specific DNA locations^12, 13^. Our results highlight that, for most TFs, the presence or absence of a sequence motif cannot reliably predict factor binding in vivo. Comparison of our ChEC data and published ChIP datasets for 102 TFs revealed that, for nearly all factors, ChEC identified more binding targets, and that ∼38% of binding targets captured by ChIP-exo are also observed by ChEC-seq (**Fig S2A-B**). For further comparison, we used ChIP-seq to map binding sites for the activators Gal4 and Gcn4 and found that ChEC identified 40-72% of the binding sites identified by ChIP respectively (**Fig S2C**).

### Promoter classification based on TF binding patterns

To categorize promoters based on TF binding events, we used unsupervised K-means clustering, resulting in separation of genes into three distinct classes based on the number of TFs bound to their promoters (**Fig 1B; Table S3A**). Cluster I includes approximately 53% of the total genes analyzed and whose promoters are bound by a relatively small number of TFs (median of 7 TFs/promoter). Cluster II promoters (∼26% of genes) are bound by a moderate number of TFs (median of 24/promoter), whereas Cluster III promoters (∼14% of genes) are bound by many TFs (median of 67/promoter) (**Fig 1C**). The ChEC signal strength of Cluster I genes (likely related to TF occupancy) is modestly higher compared to signals at Cluster I and II genes (**Fig S3A, Table S3B**). Cluster I and Cluster II are enriched in TFIID class genes whereas Cluster III is enriched in CR class genes (**Fig 1D**). While Cluster III promoters have wider nucleosome depleted regions (NDRs) compared to the other two clusters (**Fig 1E**; median of 250 bp vs 125-140 bp), other promoter features likely play major roles in the large difference in the number of TFs that interact with these regulatory regions. Since the in vivo DNA binding behavior of many TFs is dynamic with lifetimes on the order of seconds^38–40^, we envision that only a fraction of these TFs bind to a single promoter at any one time. The expression levels of Cluster I genes are, on average, relatively low, while Cluster III genes include many highly expressed genes (**Fig S3B**). Moreover, we observed overlap of our clusters with prior gene classes based on ChIP-exo data^30^ : Cluster I is enriched for UNB class genes, Cluster II for TFO genes, and Cluster III for STM genes (**Fig S3C**). This earlier study suggested that many yeast genes (∼46% - UNB genes) lacked detectable binding of any TF to their promoter region while the TFO promoters (∼33%) interact mostly with nucleosome positioning factors. Our results are in contrast with those conclusions, as 5425 promoters are bound by at least one sequence specific TF and these differences are likely due in part to the increased sensitivity of ChEC-seq in detecting TF-DNA binding compared with ChIP-exo.

To mine our binding data for TFs that tend to occupy the same set of promoters, we performed hierarchical clustering on the correlation of TF binding patterns (**Table S5A**). This revealed TF binding clusters that show remarkable agreement with previously established TF-TF interactions along with many new examples (**Fig 1F**). For example, interactions identified by our analysis that agree with prior findings include Gal4/80, Med8/15, Hap2/3/5, Hir2/3, NuA4 subunits, Snf5/Swp82, Tup1/Cyc8, Kar4/Dig2/Tec1, Stp1/2, Rtg1/3, Bur6/Mot1 and MBP/SBP complex proteins. Further, we observed that TFs with similar known function fall into the same cluster, such as Fhl1/Ifh1 (ribosomal protein (RP) gene regulation) Cha4/Aro80 (amino acid catabolism), Ert1/Rds2 (gluconeogenesis), and Skn7/Hms2 (heat stress).

Additionally, we identified a cluster of TFs including subunits of the NuA4 HAT transcription cofactor complex (**Fig 1F**; top left corner highlighted in black) that are enriched at Cluster II genes, consistent with findings that NuA4 complex works predominantly at TFIID-dependent genes ^24, 41^. Another cluster contained factors enriched at Cluster III promoters including subunits of the cofactor Mediator (Med8/15) which is consistent with Med Tail dependence at a subset of CR promoters ^25^ (**Fig 1F**; mid right cluster highlighted in black). These results further validate our binding data and provide a rich source for exploring previously unknown TF binding interactions.

### Activation domain-mediated binding at Cluster-III promoters

Our findings suggest that the DBD sequence preferences of most yeast TFs can explain only a modest fraction of genome-wide binding. Consistent with this, recent studies showed that the non-DBDs of many yeast TFs are important for DNA targeting^12, 13^. To determine if TF targeting via activation domains (AD) might explain our results, we created mutations in Gcn4, a well-characterized strong activator involved in the response to amino acid starvation. Gcn4 mutations were designed to inhibit either DBD or AD function. The Gcn4-dbd mutant contains a triple mutation in residues (N235A, R243A, S242A) ^42–44^ important for DNA binding. The Gcn4-ad mutant has a deletion of both of the Gcn4 ADs (Δ2-134) ^45, 46^. Western analysis showed that both mutants were expressed at a higher level compared with WT Gcn4 (**Fig S4A**). ChEC-seq mapping of Gcn4-dbd showed a loss of Gcn4 binding mostly at sites containing the Gcn4 consensus motif (**Fig S4 B, C**). Conversely, binding of Gcn4-ad was reduced at ∼75% of the sites lacking a consensus motif. However, Gcn4-ad binding was surprisingly also reduced at about half of the motif-containing sites (**Fig S4C**), suggesting that the Gcn4 AD plays an important role in targeting and/or stabilizing many Gcn4-DNA interactions whether or not they are driven by DBD-motif interactions. Overall, Gcn4-DNA binding exhibits three distinct modes: DBD (29%), AD (31%) or AD + DBD-mediated binding (36%).

Strikingly, we observed that Gcn4 DBD-independent binding is greatly enriched at Cluster III promoters and, consistent with this observation, the Gcn4-ad mutant shows a dramatic loss in binding at Cluster III promoters (**Fig S4D**). Further analysis showed that the Gcn4 DBD-independent binding pattern showed good correlation with SWI/SNF subunits, Swi3 and Swp82 (Pearson’s correlation r = 0.5), Mediator subunit, Med15 (r = 0.5) and with negative regulators of transcription Bur6, Mot1 (r = 0.6 and 0.5 respectively) (**Fig S4D**). Apart from wider NDRs, Cluster III genes are enriched for genes that reside near boundaries of chromosome interaction domains^47^ (**Fig S4D**; right column). Consistent with these results, Mediator and chromatin remodelers have a role in higher order chromatin structure and are known to be enriched at boundary elements^48, 49^. Our findings highlight the importance of non-traditional binding and targeting mechanisms in genome-wide TF function.

### Functional targets of sequence specific transcription factors

If one imagines that TFs function equivalently at all bound genes, the number of bound promoters is surprisingly large for many of the yeast TFs. For example, Ste12, involved in mating and pseudohyphal/invasive growth pathways, is found at >3000 promoters yet prior studies suggest that Ste12 regulates only a small number of genes. To systematically evaluate the genome-wide functions of TFs and to determine the degree to which binding dictates function, we assayed changes in nascent RNA levels following rapid depletion of TFs using the auxin induced degron (AID) system^50^. A set of 135 yeast strains were created containing the AID tag fused to nearly all the sequence-specific TFs for which we obtained clear ChEC binding results (**Fig 2A**). Unless otherwise indicated, cells were grown in synthetic complete glucose media. Following 3-Indole Acetic Acid (IAA) treatment, cells were treated with 4-thiouracil (4TU) to label newly transcribed RNAs, followed by nascent RNA enrichment, RNA-seq^51^ and Western blot analysis (**Fig 2B**). Most of the TF-AID fusions showed dramatic reduction in protein levels after 30 min of hormone treatment (representative examples shown in **Fig S5A**). Because some earlier studies suggested that many TFs only modestly contribute to transcription of individual genes^19, 23^, we used a lenient threshold to score expression changes (padj < 0.1 and fold change > ± 1.3) and defined TF expression target genes as those exceeding this threshold. Data was collected in triplicate and, for most of the strains, there was < 20% average variation in nascent mRNA levels between replicates **(Table S4**).

**Figure 2.**
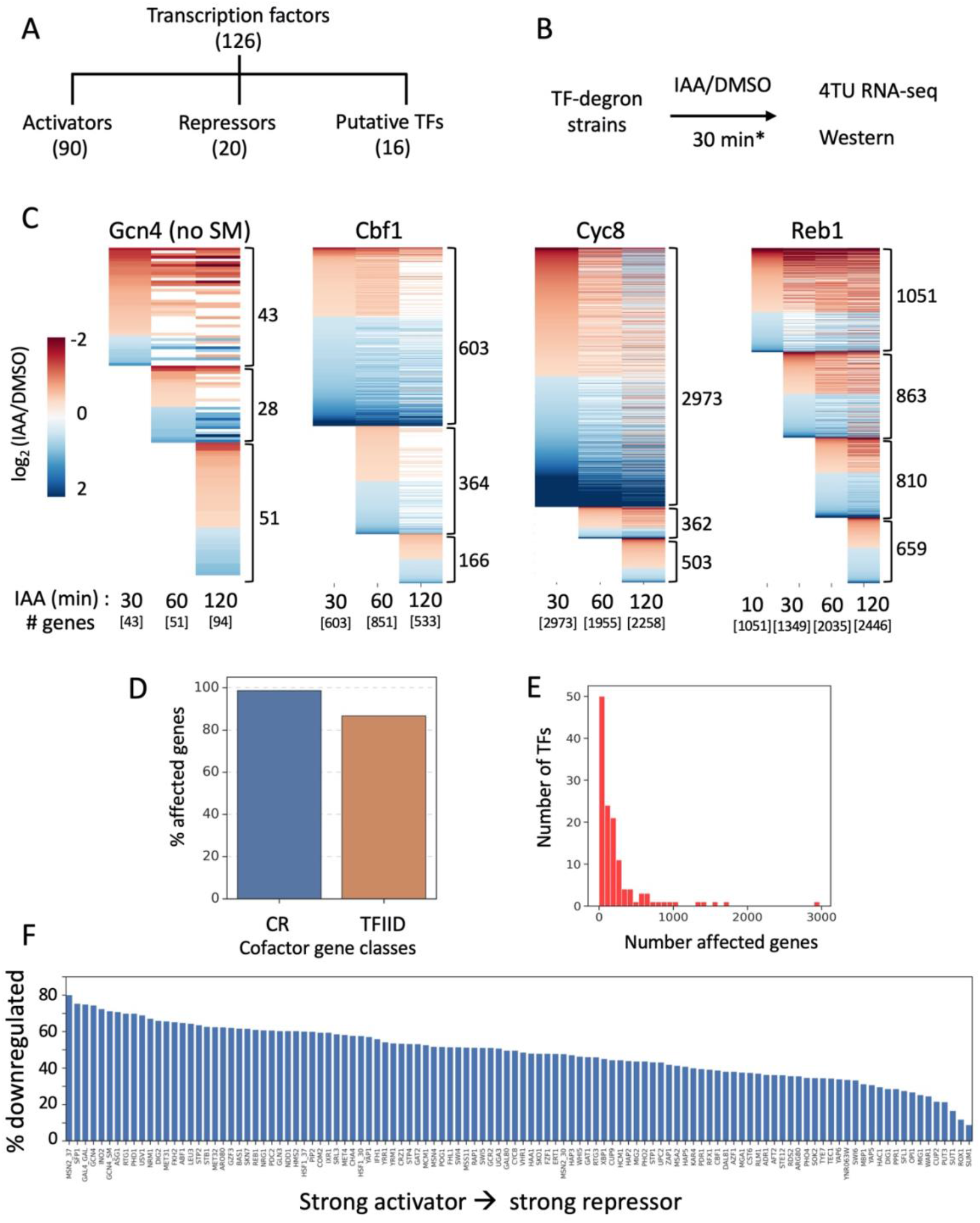
Identification of TF expression targets by rapid depletion. **(A)** Schematic showing functional categories of 126 TFs analyzed by depletion and 4TU RNA-seq analysis. Categories are assigned based on their description in the SGD database^55^. **(B)** Outline of TF depletion strategy. The set of TF-degron strains (in triplicate) was treated with either IAA or DMSO for 30 min (*except conditions noted in panel C) followed by 5 min of RNA labeling with 4-ThioU and subsequent analysis by RNA-seq and Western blot. **(C)** Heatmaps display time course of log2 fold changes in transcription upon rapid depletion of Gcn4, Cbf1, Cyc8, or Reb1. Differentially expressed genes were identified using the DESeq2 package^56^ with a threshold padj < 0.1 and fold change greater than ± 1.3. The color bar ranges from −2 to +2 log2 fold change. Total number of affected genes at each time point are shown in brackets below and the number of additional genes affected at each timepoint are shown on the side in brackets. Note that some genes affected at earlier time points are recovered at later time points (see text). **(D)** Plot shows the distribution of expression targets among the cofactor gene classes (promoter classification^26^ is defined for 4322 total protein coding genes among 4987 affected genes). The x-axis displays percent of TFIID and CR classes, and the y-axis shows the percentage of genes in each class affected by depletion of one or more TFs. **(E)** Histogram displays the frequency distribution of expression targets for 126 TFs determined after 30 min of IAA treatment. The x-axis shows the number of expression targets, while the y-axis displays the number of TFs. Using our criteria for expression changes, a total of 4987 genes are affected upon depletion of at least one TF. **(F)** Most yeast TFs have dual activation and repressor functions. Transcription factors ranked by activator function as measured by % of regulated genes that are downregulated upon rapid depletion.

We first investigated the optimum time for TF depletion with the goal of maximizing direct but minimizing indirect effects on transcription. RNA was harvested at 3-4 time points following Gcn4, Cbf1, Reb1, or Cyc8 depletion (**Fig 2C**). 30 min after IAA addition to the Gcn4-degron strain (in the absence of amino acid starvation), 43 genes showed altered expression. However, after prolonged Gcn4 depletion (60-120 min), some of the previously affected genes recovered (20, 18 respectively), and expression from additional genes (28, 51 respectively) were affected, suggesting secondary effects (**Fig 2C**). Likewise, after depleting Cbf1 for 30 minutes, expression from 603 genes were initially affected, but additional genes (364 and 166) showed expression changes at 60 and 120 minutes of depletion, while about half (316) of the initially affected genes recovered at 120 minutes. Strikingly, after depleting Cyc8 for 30 minutes, transcription from nearly half of protein coding genes (2973) was altered. However, after 60 minutes, 1380 of these genes recovered while expression from 362 additional genes was affected. Interestingly, some of the differentially expressed genes (456) at 30 min exhibited a reverse trend at 120-minutes of depletion. For Reb1, we observed differentially expressed genes as early as 10 min after IAA addition. Overall, we observed an increase in the number of affected genes with increasing Reb1 depletion time. As Reb1 is known to be involved in NDR formation ^52, 53^, its prolonged absence may result in a ripple effect of chromatin changes spreading to neighboring genes. Reb1 can also act as a chromatin insulator, preventing the spread of negative or positive influences on neighboring promoters^54^. Consequently, the effects observed on neighboring genes without a Reb1 binding site might also be attributed to the loss of insulator function. Consistent with these hypotheses, many genes affected upon Reb1 depletion were within 10 kb but not proximal to its binding sites, and expression changes in the number of distant genes increased with longer depletion times (**Fig S5B**). This observation indicates that some TFs can impact gene expression without directly binding to the promoters of their expression targets – a point observed repeatedly for other TFs (see below).

After considering the above results, TF depletion for 30 minutes was selected as optimal for detecting maximum TF targets and minimizing secondary effects. For all subsequent experiments, transcription changes in nascent mRNA were quantitated 30 min after IAA addition to identify TF regulatory targets. After testing 135 TF-degron strains, we obtained reliable data from 126 TF-degrons that had both reproducible RNA-seq data and efficient TF depletion. Using this TF-degron set, expression from 4987 genes were altered by depleting at least one TF and, on average, one gene is regulated by a median of 3 TFs but ranges from 1-119 regulatory TFs/gene. This list includes nearly all TFIID-dependent (87%) and CR class genes (99%) (**Fig 2D**), and we observed that a larger median number of TFs influence CR genes (10 TFs) than TFIID class genes (3 TFs) (**Fig S5C**). Similarly, Cluster-III genes are regulated by a higher number of TFs when compared to the other two clusters (**Fig S5D**). Despite being regulated by a relatively larger number of transcription factors (TFs), the median impact in terms of fold change on Cluster-III genes is comparable to the other two clusters (**Fig S6A**, **S6B**). The distribution frequency of affected genes suggests that the majority of TFs regulate less than 200 genes, and a typical TF regulates 112 genes but ranges from 1-2973 genes/TF (**Fig 2E**). Surprisingly, there are very few TF that are strongly biased for either activator or repressor function **(Fig 2F**). We found that only ∼10% of TFs fit the criteria of strong activators (≥70% of total affected genes are downregulated after TF depletion) while ∼ 15% of TFs are strong repressors. In contrast, most yeast TFs have dual function with numerous genes up or down regulated after depletion. This shows that strong activators or strong repressors (e.g., Gal4 or Rox1) are atypical compared with most TFs.

### Gene clusters defined by response to TF depletion

We used K-means clustering to identify clusters of genes that share similar TF dependence patterns. Only genes that are significantly affected by individual TF depletion had their log2 fold change values recorded while a value of zero was entered for genes that did not meet the significance threshold (**Table S3C**). This analysis segregated genes into 9 unique groups (1-9) and GO enrichment analysis was used to characterize each group (**Fig 3A**). Group-1 contains largest gene set (2275), encompassing a broad array of cellular processes, and these genes are not strongly enriched for either the TFIID or CR classes (**Fig 3B**). In contrast, group-7 (carbohydrate metabolism) is strongly enriched for CR vs TFIID genes (24% CR vs 5% TFIID; hypergeometric test p-value = 4.20E-49) whereas groups 3 and 6 (involved in gene expression and cell processes) are enriched for TFIID genes (p-value = 9.57E-13 and 1.55E-10 respectively). Most of the RP genes lie in group-8 (**Fig 3C**) and genes involved in ribosome biogenesis are mostly concentrated in group-2. We observed clusters that are specific to individual TFs like Abf1 (group-3), Reb1 (group-6) or Cyc8 (groups-2 and −7). We also identified two clusters containing genes that are affected by most TFs depletions: Group-9 genes, most of which are downregulated upon TF depletion, are enriched for a role in ion transport whereas group-5 genes, most of which are upregulated, are enriched for genes involved in oxidative stress. To determine whether any genes show transcription changes due to auxin treatment rather than TF depletion, we treated an otherwise isogenic strain lacking a degron with IAA for 30 min and analyzed by 4TU RNA-seq. A total of 51 genes (modestly enriched in groups-5 and -9) were affected, suggesting that these genes are not direct targets of many TFs (**leftmost lane in Fig 3A** labeled WT).

**Figure 3.**
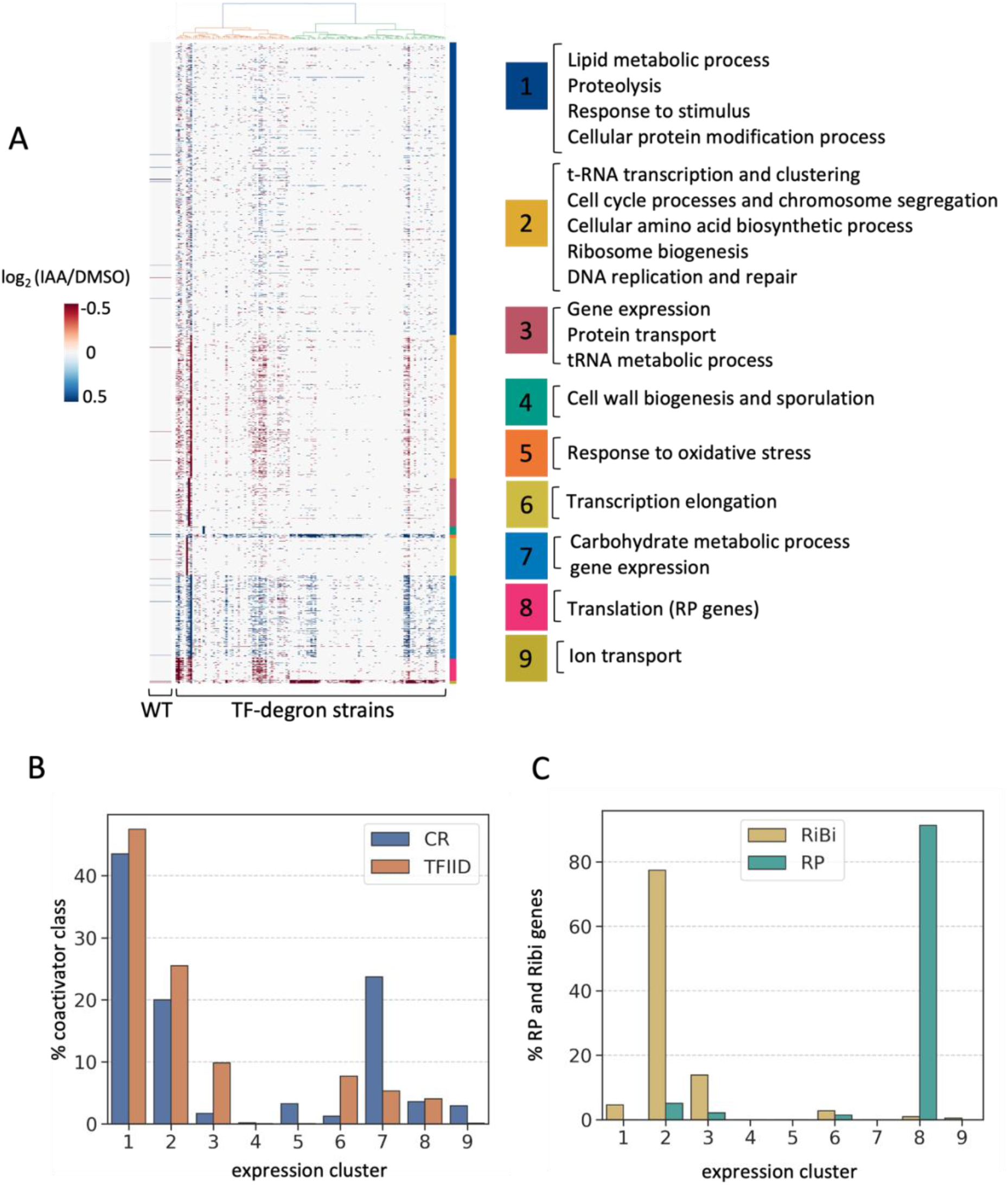
Gene clusters that respond similarly to TF depletion. (A) An unsupervised K-means clustering of expression data is represented in a heatmap, displaying 9 clusters. The y-axis denotes log2 fold change values (IAA/DMSO) for 4987 genes following depletion of TFs relative to DMSO addition, and the x-axis shows data for 126 TFs. The column labeled “WT” displays genes that are altered in expression after addition of IAA to the parent strain lacking a degron. Enriched GO terms of biological process obtained for each group 1-9 using Panther^57^ are shown on the right. (B) The plot displays the distribution of cofactor dependent gene classes in groups 1-9. The y-axis shows the percent of expression targets in each coactivator class (TFIID in red and CR in blue), and the x-axis represents the clusters derived in 3A. (C) Plot shows the percent distribution of ribosomal protein (RP) and ribosome biogenesis (RiBi) genes in the derived groups. Group-8 contains mostly RP genes while RiBi genes are concentrated in group-2.

Further validating our findings, we observed good correlation in expression targets for sets of TFs known to function together. For example, genes affected by depletion of Rap1 and Ifh1 show good correlation (r = 0.54; **Table S5B**). Likewise, 72% of Rox1 gene targets overlap with Cyc8, that are involved in ergosterol biosynthesis. As expected, depletion of TFs from the same complex are often well correlated, such as Rtg1 and Rtg2; Met4, Met31 and Met32; Hap3 and Hap5 (**Table S5B**). Finally, we observed that five transcription factors Abf1, Rap1, Reb1, Swi4 and Cyc8 target the largest number of genes (1567, 1689, 1349, 1434, 2973 genes respectively), demonstrating that they play the most general roles in yeast gene expression under the tested growth conditions. Another 11 TFs (Gat2, Gcr2, Leu3, Ifh1, Met4, Hcm1, Msa1, Met31, Cbf1, Stp1, Nrg1) target between 1004-561 genes, while most other TFs regulate many fewer genes, emphasizing their more specialized roles.

### A surprising relationship between TF binding and expression targets for most TFs

To investigate the overlap between TF binding and transcriptional regulation we integrated the results from our TF binding and depletion studies. Surprisingly, we observed moderate overlap (>40%) between TF binding and expression targets for only a few TFs (e.g., Ifh1, Cyc8, Rap1, Abf1 and Reb1), where the TF both binds to the gene upstream regulatory region and expression of the nearby gene responds to TF depletion (**Fig 4A**). In contrast, for most TFs, only a small subset of promoters with binding targets have altered expression upon depletion of those TFs. We obtained similar results when comparing TF binding mapped by ChIP-exo^30^ with our expression data, showing that the poor correspondence between binding and transcriptional response for many TFs is not specific to the mapping technique (**Fig S7A**). Some of the apparent non-functional TF binding may be due to redundancy with other factors or condition-specific function of the TFs (see below). In any event, our findings show that for most TFs, binding is often a poor predictor of whether expression of that gene responds to the bound TF.

**Figure 4.**
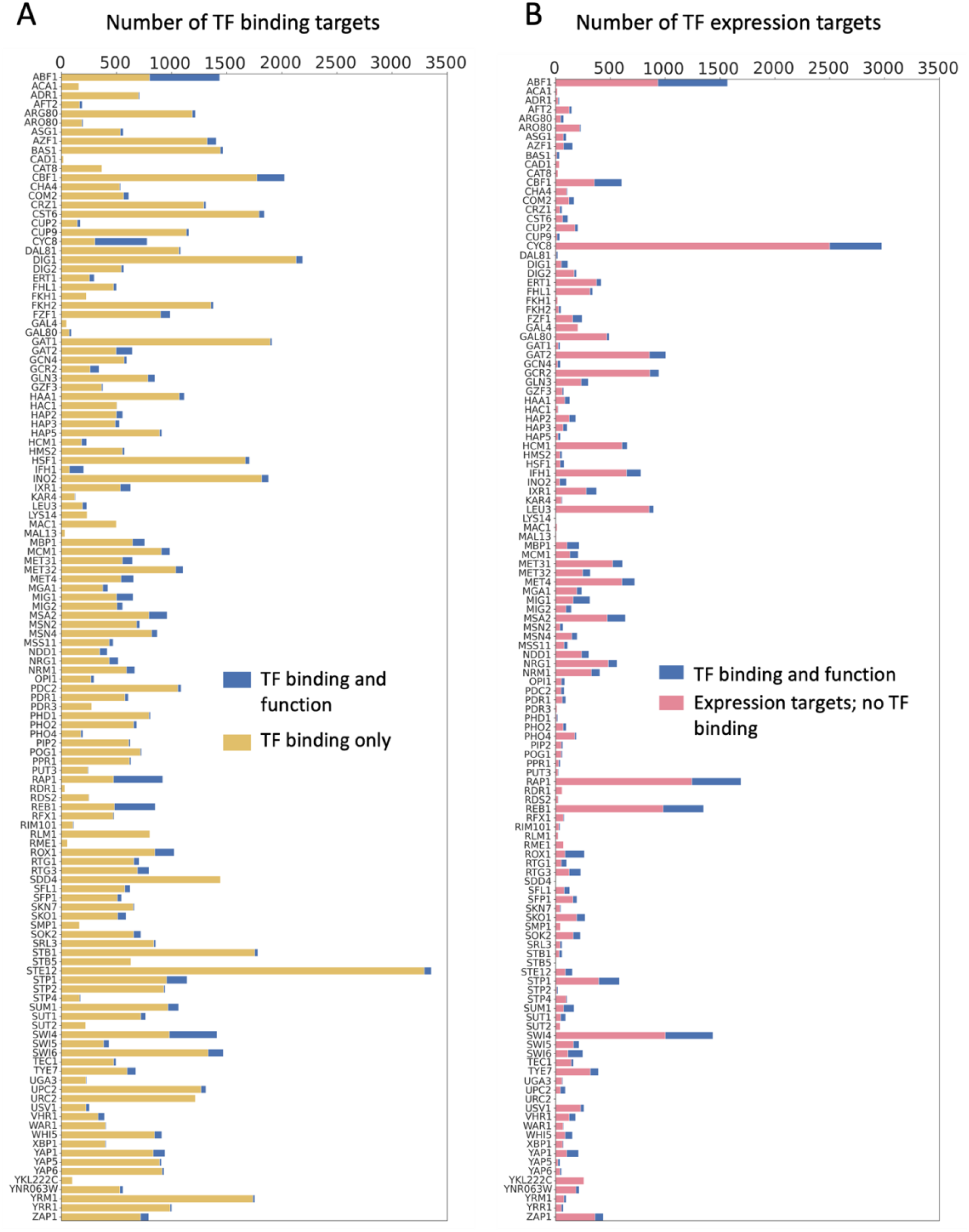
Integration of TF binding and expression data. **(A)** The bar plot displays the fraction of promoter proximal and functional TF binding sites among the total number of binding sites. The y-axis represents each TF and x-axis displays the number of binding targets detected by ChEC-seq. The blue bar indicates functional binding targets, while the yellow bar displays binding targets in gene regulatory regions where transcription of the downstream gene is not altered by factor depletion. **(B)** The bar plot displays the proportion of functional binding sites in relation to the total number of expression targets. The y-axis represents each TF and x-axis displays the number of expression targets. The blue bars indicate the number of functional targets where the TF binding site lies in the regulatory region of the affected gene, and the pink bars represent expression targets without detectable TF binding measured by ChEC-seq.

### Many expression targets do not contain a TF binding site in their gene regulatory region

Another striking observation for many TFs was that most of the expression targets revealed by the 30- minute TF depletion lacked detectable binding for that TF in the upstream gene regulatory region (**Fig 4B**). For example, the general transcription repressor Cyc8 detectably binds 777 promoters, but a 30- minute depletion affects expression from 2973 genes (474 bound by Cyc8). Likewise, the general regulatory factor Reb1 binds 851 promoters, but depletion alters expression of 1349 genes (369 bound by Reb1). This discrepancy is not due to mapping TF binding using ChEC-seq since a similar difference between TF binding and expression targets is observed using ChIP-exo binding data (**Fig S7B**). The strategy of 30 min TF depletion coupled with measurement of changes in newly synthesized RNA is designed to minimize secondary affects but it’s possible that some of the affected genes may be secondary TF targets. However, as described above, we also examined transcription changes after only 10 min of Reb1 depletion (**Fig 2C**) and again found a large discrepancy between TF binding and functional targets; 10 min after IAA addition to the Reb1-degron strain, expression from 1051 genes was altered but only 278 of these gene promoters are bound by Reb1. This result again shows that it is common for yeast TFs to regulate genes in the absence of detectable binding to the affected promoters. Since both ChEC and ChIP-exo failed to detect TF binding at many expression targets, as an alternative approach, we used the presence of a TF motif as a proxy for binding. We used FIMO^36^ to scan for TF motifs in the gene regulatory regions (400 bps upstream and 200 bps downstream of TSS) of expression targets and found that the majority of expression targets lack a motif (**Fig S8A**), in agreement with our conclusions above.

Our expression analysis shows that (i) ∼8% of TF binding targets are functional under the assayed growth conditions and (ii) only ∼24% of expression targets identified by RNA-seq show detectable binding for the corresponding TF (**Fig S8B**). To determine whether the observed overlap between TF binding and expression response was due to random chance or if expression targets are enriched for TF binding, we employed a hypergeometric test, using binding data from either ChEC-seq or ChIP-exo (**Fig S8C, D**). This analysis showed that there is significant overlap between TF binding and expression targets for 73% and 65% of the TFs assayed by ChEC and ChIP-exo respectively. In other words, although many expression targets lack detectable TF binding, for 73% of the tested TFs, the observed overlap between binding and function is not by chance given the number of TF-bound, differentially expressed, and total analyzed genes (5891).

### Characterization of functional TF binding sites

For the following analysis, we defined promoters that are both bound and regulated by a TF as functional binding targets. 116 TFs out of 126, have at least one common binding and expression target. Based on the frequency distribution (**Fig 5A**), a yeast TF typically has 32 functional binding targets, with most TFs having less than 50 and a few showing > 200. While expression from nearly all genes is affected by rapid depletion of one or more TFs (**Fig 2E**), out of the 4987 genes affected, only 2517 genes are both bound and regulated by at least one TF (**Table S3D**). This latter gene set includes 39% of the TFIID class genes (1652 genes) and 84% of the CR genes (557 genes) (**Fig 5B**). Since the CR genes are biased for functional TF binding targets compared with TFIID genes, we analyzed our data to determine the fraction of TF binding sites that are functional for each TF and whether this fraction differs based on the coactivator class of the target promoter. We found that, for the majority of TFs (72%), the fraction of functional binding targets is higher in the CR class genes by at least 2-fold (**Fig S8E**). This observation suggests that transcription from CR class genes is more responsive to promoter-bound TFs compared to TFIID genes. To extend these findings, we examined our data to identify TFs that function preferentially at TFIID or CR promoters, calculating the percentage of functional binding targets for each TF in each coactivator gene class. After filtering out TFs that have less than 20 functional targets, we found 19 TFs that contain at least 70% functional targets in the CR class (**Fig 5C**). This list includes well characterized stress responsive TFs such as Gcn4, Hsf1 and Msn2. We also identified 8 TFs with at least 70% functional targets that belong to TFIID class genes, and this list includes general transcription regulators like Rap1, Reb1 and Abf1 (**Fig 5C**). Finally, we performed hierarchical clustering using TF correlations based on fold change values of functional binding targets to determine which factors function together through binding sites in the gene regulatory regions (**Fig 5D**). This analysis is distinct from the clustering in **Fig 1**, where TFs are clustered based on binding alone without regard to regulation of the nearby gene. In agreement with known relationships, we found examples of clustered TFs with well characterized interactions, such as Rtg1/3, Pho2/4, Mbp1/Nrm1, Gal4/80, Mig1/2, Met4/31/32, and Hap3/5 as well as many more suggested functional relationships. For example, TFs that are known to regulate similar pathways such as Ifh1/Rap1, Mcm1/Arg80 also clustered, suggesting possible synergistic mechanisms.

**Figure 5.**
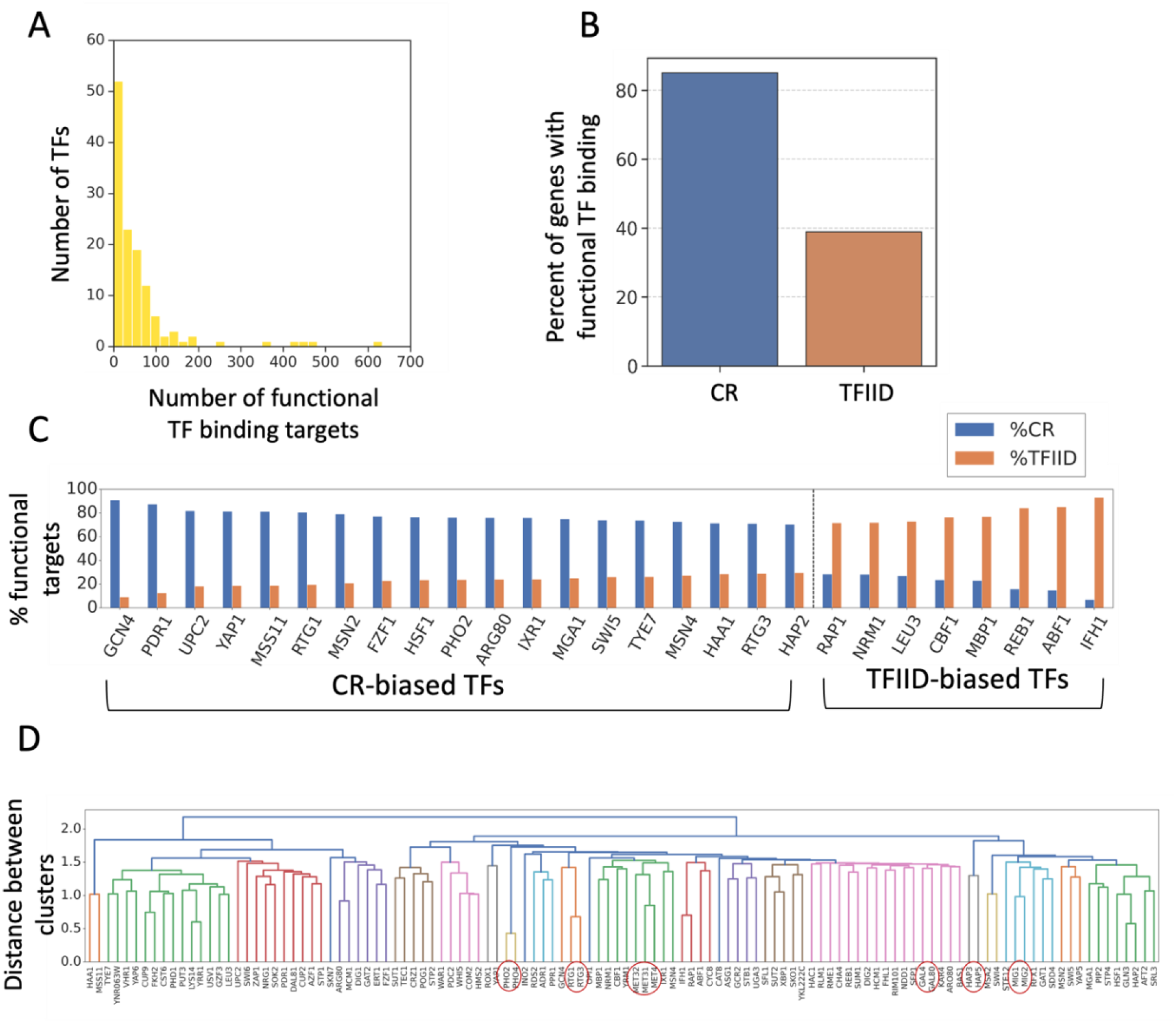
Characterization of functional TF targets. **(A)** Histogram illustrates the distribution of functional binding targets for 116 TFs (cases where the bound TF regulates the nearby gene). The x-axis represents the range of these functional targets, while the y-axis shows their frequency. **(B)** Plot displays the distribution of genes containing functional TF binding targets in the TFIID and CR coactivator class genes. The y-axis represents the percent of genes in each gene class that are affected by depletion of one or more TF and also contain a binding site for that TF in the gene regulatory region. **(C)** Bar plot showing TFs that have functional targets enriched in TFIID or CR class genes. 19 TFs (CR-biased TFs) have functional targets that are at least 70% CR while 8 TFs (TFIID-biased TFs) have functional targets that are at least 70% TFIID. **(D)** Hierarchical clustering was used to determine the correlation of functional TF binding sites based on log2 fold change values of gene expression changes upon TF depletion. Results are represented as a dendrogram. Distance between clusters is obtained using average method and shown on y-axis. Several examples of known TF interactions are highlighted by red circles.

It is surprising that only a small subset of binding targets has functional consequences, which may indicate that some transcription factor (TF) binding sites are active only under specific growth conditions. To investigate this possibility, we depleted several TFs during specific stress conditions where TF importance has been previously demonstrated. We first examined Gcn4, inducing amino acid stress by adding sulfometuron methyl (SM) for 60 min followed by Gcn4 depletion for 30 minutes. The depletion of Gcn4 under SM stress affected expression from 296 genes (**Fig S9A**), while only 43 genes were affected under non-SM stress conditions (**Fig S9A; Fig 2C**). Based on our ChEC data, we observed that 118 out of the 296 genes affected are bound by Gcn4 when treated with SM (**Fig S9E; Table S1**). Next, we subjected cells to heat shock (for 10 min) followed by depletion of Hsf1 or Msn2. Depletion of Hsf1 after heat shock led to changes in expression of 153 genes, compared to 78 genes under non-heat shock conditions (**Fig S9B**). Out of the 153 differentially expressed genes, 82 genes are bound by Hsf1 (**Fig S9E; Table S1**). Similarly, depletion of Msn2 following heat shock affected 131 genes (**Fig S9C**), of which 64 gene promoters were occupied by Msn2 (**Fig S9E**; **Table S1**). Finally, after galactose induction followed by Gal4 depletion, expression of 24 genes is affected (**Fig S9D**), of which 6 show Gal4 binding to their promoters (**Fig S9E**; **Table S1**). This contrasts with no expression changes observed upon depletion of Gal4 in the absence of galactose. Our combined results reveal a notable increase in the fraction of functional binding sites for TFs under growth conditions that specifically depend on the functionality of these TFs (**Fig S9F**). However, for this small set of TFs, our analysis revealed that majority of TF binding sites had no apparent role even under conditions known to activate TF function.

## Discussion

Much remains to be learned about how TFs locate their targets and the mechanisms by which they combine to generate functional regulatory elements and control transcription of individual genes. By integrating information on the binding and expression targets for the near-complete set of TFs from a simple eukaryote, we have gained important and surprising insights into the role of TFs and the mechanisms whereby they regulate genome-wide transcription.

With a few exceptions, many of the binding sites for most TFs lack a corresponding DNA sequence motif. The widespread extent to which this non-motif binding mode is used was unexpected but is consistent with alternative modes of TF-DNA recruitment and binding, such as recognition of DNA shape or targeting mediated by disordered regions of the TFs^3, 12, 13^. In one case tested here, the Gcn4 AD modulates much of the motif-independent binding and these sites are found mainly in regulatory regions of Cluster III genes. It is striking that genes in this cluster interact on average with very large numbers of TFs, are enriched for highly regulated genes, and that many of these genes are located near the boundaries of chromosome interaction domains. We speculate that TF clustering and/or higher order chromatin structure may contribute to the unusually high numbers of TFs found at these genes.

Comparison of TF binding and expression targets revealed an unexpected and complex relationship between TF binding and transcriptional regulation. The conventional model for TF function, that has been validated over many years at individual genes, is that nearly all yeast TFs function from binding at UAS elements, located a short distance upstream from their regulatory targets. While our data finds many instances of this mechanism (functional TF binding sites), it is very surprising that a much greater number of instances do not fit this model. We have found a great many examples where rapid TF depletion affects gene expression where there is no detectable binding of that TF to the upstream region of the affected gene. While some fraction of these instances may be due to indirect effects, we find that regulatory targets without an upstream TF binding site are apparent within as little as 10 min after initiating TF depletion. Our data suggests that this is a widespread phenomenon. For example, while nearly all genes are regulated by at least one TF and have at least one TF bound to the regulatory region, only about half (2517) of the 4987 genes affected have a functional TF binding site. Our results also contradict a recent model that proposed most yeast genes that are constitutively expressed have a common architecture and do not respond to regulatory TFs^30, 58^. Here we demonstrate that nearly all Pol II transcribed genes respond to the loss of individual TFs and that the magnitude of the response is roughly similar for all classes of genes tested.

Finally, we found that, for most TFs, only a fraction of bound TFs regulates the nearby gene, showing that the binding of a TF does not automatically correspond to regulation of the linked gene. Part of this might be explained by the widespread repression of transcription by Cyc8; we found that over a quarter of the protein coding genes are derepressed upon rapid Cyc8 depletion and this repression may prevent the function of many TFs. It’s important to note that nearly all our measurements were carried out in glucose-grown, non-stressed cells. As such, it is expected that many stress-activated pathways are repressed with corresponding repression or inactivation of stress-induced TFs. Consistent with this, assaying the binding and expression targets under several stress conditions led to a higher fraction of functional TF binding. However, even under these activated conditions, the fraction of non-functional TF binding is still >80%, suggesting that our conclusions are not strictly dependent on the growth conditions used for most of our assays. The combination of findings reported here provide a deeper understanding of TF-mediated gene regulation and provide a rich resource to delve deeper into mechanisms connecting the dynamic interplay between TFs and their effect on gene expression.

## Methods

### Yeast strains and cell growth

Yeast strains are listed in **Table S6**, and *S. cerevisiae* strains are derivatives of BY4705^59^. Most degron and MNase-tagged strains were constructed as previously described^25, 26, 60^ . However, in some instances, tagging proteins with the full-length IAA7 degron led to a growth phenotype. In these cases, proteins were tagged with a shortened IAA7 degron^24, 25^ that resulted in little or no growth phenotypes for strains used in this work. When either of these approaches led to growth phenotypes, N-terminal degron tagging was used. Some strains were constructed using conventional homologous recombination (SHY1667, 1668, 1669, 1670, 1671, 1672, 1698, 1699), while other strains (SHY1704, SHY1705, and

SHY1706) were constructed using CRISPR/CAS9-mediated gene editing. For N-terminal tagging, proteins were tagged with a shortened IAA7 degron (termed mini degron; 13 kDa total) that was inserted just after the ATG in the coding region. The sequence of the mini degron tag is below. Small letters indicate the 3X V5 epitope tag, bold letters are linker sequence, and non-bold capital letters encode the shortened IAA7 degron:

**GCT**ggtaaacctatacctaatccattattgggactagatggaaaaccaataccaaatcccttacttggtttggattctacaccaattcctaatcctctattaggactggatagtaca**GGTGCCGGTGCTGGTGCCGGAGCTGGCGCAGGTGCT**AAGGAAAAATCTGCGTGTCCAAAGGACCCTGCAAAACCACCAGCCAAGGCACAAGTT GTAGGTTGGCCCCCTGTAAGATCCTATAGAAAGAATGTTATGGTTTCTTGCCAAAAATCTTCTGGAGGCCCTGAAGCAGCTGCATTTGTTAAAGTTAGTATGGACGGTGCTCCTTACTTGAGAAAAATAGACTTGAGAATGTATAAA

PCR was used to amplify the mini degron tags and primers included 50-53 bps of gene-specific homology for homology directed repair. Primers for ABF1 amplification also included a G to A mutation in the PAM sequence immediately 5’ of the ATG. Guide sequences were cloned into pCAS (Addgene 60847) using Quickchange and the primer 5’-CGGGTGGCGAATGGGACTTT [20-mer guide sequence] GTTTTAGAGCTAGAAATAGC-3’, plus its reverse complement. Guide sequences for each factor are as follows: ABF1 – 5’-ACCATATTTGCAATTTCACA-3’; GAL4 – 5’-CAGTAGCTTCATCTTTCAGG-3’; MCM1 – 5’- AAAATGTCAGACATCGAAGA-3’. Resulting pCAS plasmids (pLH411, ABF1; pLH412, GAL4; pLH413, MCM1) and each gene-specific mini degron PCR amplicon were transformed into yeast strain SHY1235 using standard yeast methods. Transformants were plated onto YPD plates, then replica printed onto YPD + G418. Single G418-resistant colonies were isolated, patched onto YPD and screened for mini degron insertion using yeast colony PCR, then sequenced and screened for loss of the pCAS plasmid.

*S. cerevisiae* strains were grown as indicated in synthetic complete (SC) media (per liter: 1.7 g yeast nitrogen base without ammonium sulfate or amino acids (BD Difco), 5 g ammonium sulfate, 40 μg/ml adenine sulfate, 0.6 g amino acid dropout mix and supplemented with two micrograms/ml uracil and 0.01% other amino acids to complement auxotrophic markers). Standard amino acid dropout mix contains 2 g each of Tyr, Ser, Val, Ile, Phe, Asp, Pro and 4 g each of Arg and Thr. *S. pombe* strains were grown in YE media (0.5% yeast extract, 3% glucose). Where indicated, *S. cerevisiae* strains at an A600 of ∼1.0 were treated with 500 μM indole-3-acetic acid (IAA) dissolved in DMSO (or with DMSO alone) for 30 min. Where indicated, cells were grown in SC (-Ile-Val) media and exposed to stress conditions by incubation with 0.5 μg/ml sulfometuron methyl (SM) dissolved in DMSO or with DMSO alone for 60 min; by addition of an equal volume of 44°C media to the 30°C culture followed by incubation at 37°C for 10 min for heat shock; or incubation for 10 min at 30°C for no heat shock; cells were grown in YEP-Raffinose (1% yeast extract, 2% peptone, 2% raffinose) and induced with 2% galactose for 2hr, prior to RNA labeling.

### ChEC-seq experiments

ChEC-seq was performed as previously described^26, 33^. *S. cerevisiae* 50 ml cultures were pelleted at 2000 x g for 3 min. Cells were resuspended in 1 ml of Buffer A (15 mM Tris, 7.5, 80 mM KCl, 0.1 mM EGTA, 0.2 mM spermine (Millipore Sigma #S3256), 0.3 mM spermidine (Millipore Sigma #85558), protease inhibitors (Millipore Sigma #04693159001)), transferred to a 1.5 ml tube and pelleted at 1500 x g for 30 sec. Cells were washed twice with 1 ml of Buffer A and finally resuspended in 570 μl of Buffer A. 30 μl 2% digitonin (Millipore Sigma #300410) was added to a final concentration of 0.1% and cells were permeabilized for 5 min at 30°C with shaking (900 rpm). 0.2 mM CaCl_2_ was added to the samples followed by incubation for another 5 min at 30°C. 100 μl cell suspension was mixed with 100 μl Stop Solution (400 mM NaCl, 20 mM EDTA, 4 mM EGTA). Stop Solution was supplemented with 5ng MNase digested *D. melanogaster* chromatin. Samples were incubated with 0.4 mg/ml Proteinase K (Thermo

Fisher Scientific #AM2548) for 30 min at 55°C and the DNA was purified by phenol/CHCl_3_/isoamyl alcohol (25:24:1) extraction and ethanol precipitation. Pellets were resuspended in 30 μl 0.3 mg/ml RNase A (Thermo Fisher Scientific #EN0531) (10 mM Tris, 7.5, 1 mM EDTA, 0.3 mg/ml RNase A) and incubated for 15 min at 37°C. 60 μl of Mag-Bind reagent (Omega Biotek #M1378-01) was added and the samples were incubated for 10 min at RT. Supernatants were transferred to a new tube and the volume was adjusted to 200 μl (10 mM Tris, 8.0, 100 mM NaCl). DNA was purified again by phenol/CHCl_3_/isoamyl alcohol (25:24:1) extraction and ethanol precipitation, and resuspended in 25 μl 10 mM Tris, 8.0.

### ChIP-seq experiments

ChIP-seq experiments were performed similarly to a published method^26^. 100 ml *S. cerevisiae* or *S. pombe* cultures were crosslinked with 1% formaldehyde (Sigma-Aldrich #252549) for 20 min in the above growth conditions, followed by another 5 min treatment with 130 mM glycine. Cells were pelleted at 3000 x g for 5 min, washed with cold TBS buffer, pelleted at 2000 x g for 3 min, flash-frozen in liquid N_2_, and then stored at −80°C for further use. Cell pellets were resuspended in 300 μl Breaking Buffer (100 mM Tris, 8.0, 20% glycerol, protease inhibitors (Millipore Sigma #04693159001)). Cells were lysed using 0.4 ml zirconia/silica beads (RPI #9834) in a Mini Beadbeater-96 (BioSpec Products) for 5 min. Lysates were spun at 21K x g for 2 min. Pellets were resuspended in 1 ml FA buffer (50 mM HEPES, 7.5, 150 mM NaCl, 1 mM EDTA, 1% Triton X-100, 0.1% sodium deoxycholate, protease inhibitors (Millipore Sigma #04693159001)) and transferred to 15 ml polystyrene tubes. Samples were sonicated in a cold Bioruptor sonicator bath (Diagenode #UCD-200) at a maximum output, cycling 30 sec on, 30 sec off, for a total of 45 min. Samples were spun twice in fresh tubes at 21K x g for 15 min. Prepared chromatin was flash-frozen in liquid N_2_, and then stored at −80°C for further use.

20 μl of the chromatin sample was used to estimate DNA concentration. First, 20 μl Stop buffer (20 mM Tris, 8.0, 100 mM NaCl, 20 mM EDTA, 1% SDS) was added to samples followed by incubation at 70°C for 16-20 hrs. Samples were digested with 0.5 mg/ml RNase A (Thermo Fisher Scientific #EN0531) for 30 min at 55°C and 1 mg/ml Proteinase K for 90 min at 55°C. Sample volume was brought to 200 μl and DNA was purified by two phenol/CHCl_3_/isoamyl alcohol (25:24:1) extractions and ethanol precipitation. DNA was resuspended in 20 μl 10 mM Tris, 8.0 and the concentration was measured using Qubit HS DNA assay (Thermo Fisher Scientific #Q32851).

20 μl Protein G Dynabeads (Thermo Fisher Scientific #10003D) was used for a single immunoprecipitation. Beads were first washed three times with 500 μl PBST buffer (PBS buffer supplemented with 0.1% Tween 20) for 3 min with gentle rotation. Beads were resuspended in a final volume of 20 μl containing PBST buffer and the indicated antibody. The bead suspension was incubated for 60 min with shaking (1400 rpm) at RT, washed with 500 μl PBST buffer and 500 μl FA buffer. Beads were resuspended in 25 μl FA buffer. 1.5 μg *S. cerevisiae* chromatin and 30 ng *S. pombe* chromatin (strain Sphc821) were combined and samples were brought to a final volume of 500 μl. 25 μl of each sample was mixed with 25 μl Stop buffer and set aside (input sample). 25 μl of beads was added to remaining 475 μl of samples followed by incubation for 16-20 hrs at 4°C.

The beads were washed for 3 min with gentle rotation with the following: 3 times with 500 μl FA buffer, 2 times with FA-HS buffer (50 mM HEPES, 7.5, 500 mM NaCl, 1 mM EDTA, 1% Triton X-100, 0.1% sodium deoxycholate), once with 500 μl RIPA buffer (10 mM Tris, 8.0, 0.25 M LiCl, 0.5% NP-40, 1 mM EDTA, 0.5% sodium deoxycholate). DNA was eluted from beads with 25 μl Stop buffer at 75°C for 10 min. Elution was repeated, eluates were combined and incubated at 70°C for 16-20 hrs together with input samples collected earlier. Samples were digested with 0.5 mg/ml RNase A (Thermo Fisher Scientific #EN0531) for 30 min at 55°C and 1 mg/ml Proteinase K for 2 hrs at 55°C. Sample volume was brought to 200 μl and DNA was purified by two phenol/CHCl_3_/isoamyl alcohol (25:24:1) extractions and ethanol precipitation. DNA was resuspended in 15 μl 10 mM Tris, 8.0 and the concentration was measured using Qubit HS DNA assay (Thermo Fisher Scientific #Q32851).

### Preparation of NGS libraries for ChEC-seq and ChIP-seq samples

NGS libraries for ChEC-seq and ChIP-seq experiments were prepared similarly as described ^26, 61^. 12 μl of ChEC samples and 5 μl of ChIP samples was used as input for library preparation. Samples were end-repaired, phosphorylated and adenylated in 50 μl reaction volume using the following final concentrations: 1X T4 DNA ligase buffer (NEB #B0202S), 0.5 mM each dNTP (Roche #KK1017), 0.25 mM ATP (NEB #P0756S), 2.5% PEG 4000, 2.5 U T4 PNK (NEB #M0201S), 0.05 U T4 DNA polymerase (Invitrogen #18005025), and 0.05 U Taq DNA polymerase (Thermo Fisher Scientific #EP0401). Reactions were incubated at 12°C 15 min, 37°C 15 min, 72°C 20 min, then put on ice and immediately used in adaptor ligation reactions. Adaptor ligation was performed in a 115 μl volume containing 6.5 nM adaptor, 1X Rapid DNA ligase buffer (Enzymatics #B101L) and 3000 U DNA ligase (Enzymatics #L6030-HC-L) and reactions were incubated at 20 deg for 15 min. Following ligation, a two-step cleanup was performed for ChEC-seq samples using 0.25x vol Mag-Bind reagent (Omega Biotek # M1378-01) in the first step and 1.1x vol in the second step. In case of ChIP-seq samples a single cleanup was performed using 0.4x vol Mag-Bind reagent. In both cases DNA was eluted with 20 μl 10 mM Tris, 8.0. Library Enrichment was performed in a 30 μl reaction volume containing 20 μl DNA from the previous step and the following final concentrations: 1X KAPA buffer (Roche #KK2502), 0.3 mM each dNTP (Roche #KK1017), 2.0 μM each P5 and P7 PCR primer, and 1 U KAPA HS HIFI polymerase (#KK2502). DNA was amplified with the following program: 98°C 45 s, [98°C 15 s, ramp to 60°C @ 3°C /s, 60°C 10 s, ramp to 98°C @ 3°C /s] 16-18x, 72°C 1 min. 18 cycles were used for library amplification for ChEC-seq samples and 16 cycles for ChIP-samples. A post-PCR cleanup was performed using 1.4x vol Mag-Bind reagent and DNA was eluted into 30 μl 10 mM Tris, 8.0. Libraries were sequenced on the Illumina HiSeq2500 platform using 25 bp paired-end reads or NextSeq 2000 using 50 bp paired-ends at the Fred Hutchinson Cancer Research Center Genomics Shared Resources facility.

### RNA labeling and RNA purification

All experiments were done in triplicates. Newly synthesized RNAs were labeled as previously described^62^. 10 ml *S. cerevisiae* or 20 ml *S. pombe* cells were labeled with 5 mM 4-thiouracil (Sigma-Aldrich) for 5 min, the cells were pelleted at 3000 x g for 2 min, flash-frozen in liquid N_2_, and then stored at −80°C until further use. *S. cerevisiae* and *S. pombe* cells were mixed in an 8:1 ratio and total RNA was extracted using the RiboPure yeast kit (Ambion, Life Technologies) using the following volumes: 480 μl lysis buffer, 48 μl 10% SDS, 480 μl phenol:CHCl_3_:isoamyl alcohol (25:24:1) per *S. cerevisiae* pellet + 50 μl *S. pombe* (from a single *S. pombe* pellet resuspended in 850 μl lysis buffer). Cells were lysed using 1.25 ml zirconia/silica beads in a Mini Beadbeater-96 (BioSpec Products) for 3 min followed by 1 min rest on ice. This bead beating cycle was repeated twice for a total of 3 times. Lysates were spun for 5 min at 16K x g, then the following volumes combined in a 5 ml tube: 400 μl supernatant, 1400 μl binding buffer, 940 μl 100% ethanol. Samples were processed through Ambion filter cartridges until all sample was loaded, then washed with 700 μl Wash Solution 1, and twice with 500 μl Wash Solution 2/3. After a final spin to remove residual ethanol, RNA was eluted with 25 μl 95°C preheated Elution Solution. The elution step was repeated, and eluates combined. RNA was then treated with DNaseI using 6 μl DNaseI buffer and 4 μl DNaseI for 30 min at 37°C, then treated with Inactivation Reagent for 5 min at RT. RNA was then biotinylated essentially as described^63, 64^ using 40 μl (∼40 μg) total RNA and 4 μg MTSEA biotin-XX (Biotium) in the following reaction: 40 μl total 4TU-labeled RNA, 20 mM HEPES, 1 mM EDTA, 4 μg MTSEA biotin-XX (80 μl 50 μg/ml diluted stock) in a 400 μl final volume. Biotinylation reactions occurred for 30 min at RT with rotation and under foil. Unreacted MTS-biotin was removed by phenol/CHCl_3_/isoamyl alcohol (25:24:1) extraction. RNA was precipitated with isopropanol and resuspended in 100 μl nuclease-free H_2_O. Biotinylated RNA was purified also as described (Duffy and Simon, 2016) using 80 μl MyOne Streptavidin C1 Dynabeads (Invitrogen) + 100 μl biotinylated RNA for 15 min at RT with rotation and under foil. Prior to use, MyOne Streptavidin beads were washed in a single batch with 3 × 3 ml H_2_O, 3 × 3 ml High Salt Wash Buffer (100 mM Tris, 7.4, 10 mM EDTA, 1 M NaCl, 0.05% Tween-20), blocked in 4 ml High Salt Wash Buffer containing 40 ng/μl glycogen for 1 hr at RT, then resuspended to the original volume in High Salt Wash Buffer. After incubation with biotinylated RNA, the beads were washed 3 × 0.8 ml High Salt Wash Buffer, then eluted into 25 μl streptavidin elution buffer (100 mM DTT, 20 mM HEPES 7.4, 1 mM EDTA, 100 mM NaCl, 0.05% Tween-20) at RT with shaking, then the elution step repeated and combined for a total of 50 μl. At this point, 10% input RNA (4 μl) was diluted into 50 μl streptavidin elution buffer and processed the same as the labeled RNA samples to determine the extent of recovery. 50 μl each input and purified RNA was adjusted to 100 μl with nuclease-free water and purified on RNeasy columns (Qiagen) using the modified protocol as described (Duffy and Simon, 2016). To each 100 μl sample, 350 μl RLT lysis buffer (supplied by the Qiagen kit and supplemented with 10 μl 1% βME per 1 ml RLT) and 250 μl 100% ethanol was added, mixed well, and applied to columns. Columns were washed with 500 μl RPE wash buffer (supplied by the Qiagen kit and supplemented with 35 μl 1% βME per 500 μl RPE), followed by a final 5 min spin at max speed. RNAs were eluted into 14 μl nuclease-free water.

### Preparation of 4TU mRNA libraries for NGS

Newly synthesized RNA isolated via 4-ThioU labeling and purification was prepared for sequencing using Ovation Universal RNA-seq System kits (Tecan) according to the manufacturer’s instructions and 50-100 ng (Universal) input RNA. rRNA was depleted during library prep using Tecan’s custom AnyDeplete probe for *S. cerevisiae* S288C. Libraries were sequenced on the Illumina HiSeq 2500 platform using 25 bp paired-ends or NextSeq 2000 using 50 bp paired-ends at the Fred Hutchinson Genomics Shared Resources facility.

### Analysis of NGS data

Data analysis was performed similarly as described^26^. The majority of the data analysis tasks except sequence alignment, read counting and peak calling (described below) were performed through interactive work in the Jupyter Notebook (https://jupyter.org) using Python programming language (https://www.python.org) and short Bash scripts. All figures were generated using Matplotlib and Seaborn libraries for Python; (https://matplotlib.org; https://seaborn.pydata.org). All code snippets and whole notebooks are available upon request.

Paired-end sequencing reads were aligned to *S. cerevisiae* reference genome (sacCer3), *S. pombe* reference genome (ASM294v2.20) or *D. melanogaster* reference genome (release 6.06) with Bowtie^65^ using optional arguments ‘-I 10 -X 700 --local --very-sensitive-local --no-unal --no-mixed --no-discordant’. Details of the analysis pipeline depending on the experimental technique used are described below.

### Analysis of ChEC-seq data

SAM files for *S. cerevisiae* data were converted to tag directories with the HOMER (http://homer.ucsd.edu)^66^ ‘makeTagDirectory’ tool. Peaks were called using HOMER ‘findPeaks’ tool with optional arguments set to ‘-o auto -C 0 L 6 F 10’, with the free MNase dataset used as a control. These settings use a default false discovery rate (0.1%) and require peaks to be enriched 10-fold over the control and 6-fold over the local background. Resulting peak files were converted to BED files using ‘pos2bed.pl’ program. For each peak, the peak summit was calculated as a mid-range between peak borders. For peak assignment to promoters the list of all annotated ORF sequences (excluding sequences classified as ‘dubious’ or ‘pseudogene’) was downloaded from the SGD website (https://www.yeastgenome.org). Data for 5891 genes were merged with TSS positions^67^. If the TSS annotation was missing (682 genes), TSS was manually assigned at position −100 bp relative to the start codon^26^. Peaks were assigned to promoters if their peak summit was in the range from −400 to +200 bp relative to TSS. In a few cases, where more than one peak was assigned to the promoter, the one closer to TSS was used. Promoters bound in at least two out of three replicate experiments were included in a final list of promoters bound by a given factor and were used to calculate promoter occupancy in all relevant experiments.

Coverage at each base pair of the *S. cerevisiae* genome was calculated as the number of reads that mapped at that position divided by the number of all *D. melanogaster* reads mapped for the sample and multiplied by 10000 (arbitrarily chosen number). Peak summits were defined using Homer as described above.

### Analysis of ChIP-seq data

For all samples coverage at each base pair of the *S. cerevisiae* genome was calculated as the number of reads that mapped at that position divided by the number of all *S. pombe* reads mapped in the sample, multiplied by the ratio of *S. pombe* to *S. cerevisiae* reads in the corresponding input sample and multiplied by 10000 (arbitrarily chosen number). The list of all annotated ORF sequences (excluding sequences classified as ‘dubious’ or ‘pseudogene’) was downloaded from the SGD website (https://www.yeastgenome.org). Data for 5891 genes were merged with TSS positions^67^. If the TSS annotation was missing the TSS was manually assigned at position −100 bp relative to the start codon. Peaks were called using HOMER as described above but with optional arguments set to ‘-o auto -C 0 L 2 F 2’, and the corresponding input dataset is used as a control.

### Analysis of RNA-seq data

SAM files for *S. cerevisiae* data were used as an input for HTseq-count^68^ with default settings. The GFF file with *S. cerevisiae* genomic features was downloaded from the Ensembl website (assembly R64-1-1). As a filtering step, we excluded all the genes that had no measurable signal in at least one out of 288 samples collected in this work under normal growth conditions. The remaining 5293 genes were used to calculate coefficient of variation (CV) to validate reproducibility between replicate experiments (**Table S4**). The log_2_ change in transcription (IAA/DMSO) and associated p-value were calculated with DESeq2^69^ using the default normalization (geometric mean is calculated for each gene across all samples, then counts for a gene in each sample is divided by mean. The median of these ratios in a sample is the size factor for that sample). The criteria used in DESeq2 to identify affected genes are a default false discovery rate (FDR) (padj < 0.1) and a minimum 1.3-fold differential expression in either direction. The effect of TF depletion on stress response was calculated by comparing the signals in corresponding IAA and DMSO treated samples (both exposed to stress)

### Motif analysis

FIMO^36^ (Finding Individual Motif Occurrences) program, from MEME-suite package is used to search set of sequences for occurrences of known motifs. Motifs in MEME format are downloaded from JASPAR database^35^ (https://jaspar.genereg.net/downloads/). FIMO uses 0-order Markov Background Model Format, therefore sequences around all 5891 TSS (400 bps upstream and 200 downstream of TSS) are used to generate 0-order background model. The optional parameters used in the command line; “--bfile --max-strand --thresh 0.00025 “

### Quantification and statistics

All NGS experiments were done in three replicates (except free MNase control done in duplicates). p-values for comparisons displayed in boxplots were determined using a Welch’s t-test and significance is defined as - ns: > 0.05, *: 0.05-0.01, **: 0.01-0.001, ***: 0.001-0.0001, ****: p <= 0.0001. Boxplots display the median value using a horizontal line inside each boxplot. To determine the significance of overlap, one-tailed hypergeometric test is employed, and an overlap is considered significant if p-value is ≤ 0.05.

### Western blot analysis

1 ml cell culture was collected and pelleted from strains after treatment with IAA or DMSO, incubated in 200 μl 0.1 M NaOH for 5 min at room temp, then resuspended in 50 μl yeast whole cell extract buffer (0.06 M Tris-HCl, pH 6.8, 10% glycerol, 2% SDS, 5% 2-mercaptoethanol, 0.0025% bromophenol blue). After heating for 5 min at 95°C, samples were centrifuged for 5 min at max speed, whole cell extracts were separated by SDS-PAGE and analyzed by Western blot using mouse monoclonal (α-V5) or rabbit polyclonal (α-Tfg2) antibodies. Protein signals were visualized by using the Odyssey CLx scanner. Uncropped Western blot data is available at Mendeley data: (https://data.mendeley.com/v1/datasets/mwcvx4f9sm/draft?a=d3058c3a-bb47-4e81-8beb-0aaf004985e5).

### Resource availability

Further information and requests for resources and reagents should be directed to and will be fulfilled by Steven Hahn.

### Materials availability

All unique/stable reagents generated in this study are available from Steven hahn without restriction.

### Data and code availability

The datasets generated during this study will be made available at Gene Expression Omnibus under accession number GEO: GSE236948.

All code snippets and whole notebooks are available from the lead contact upon request.

Any additional information required to reanalyze the data reported in this paper is available from the upon request.

## Acknowledgements

We thank Olivia Sommers for strain construction, members of the Hahn lab for comments and suggestions throughout this work, Sarah Swygert for sharing analyzed micro-C data, T. Tsukiyama, and J. Schofield for comments on the manuscript. Supported by NIH grants R35 GM140823 to SH and P30 CA015704 to the Fred Hutch Genomics and Computational Shared Resources facility.

## Author contributions

Conceptualization, L.M., S.H.; Investigation, L.M., L.W., S.H.; Formal Analysis, L.M., R.D.; Writing, L.M., L.W., S.H.; Funding Acquisition and Supervision, S.H.

---

**Figure S1. ChEC-seq mapping of yeast transcription factors (TFs)**

**(A)** Schematic showing the functional categories of 178 TFs with DNA binding successfully analyzed by ChEC-seq.

**(B)** Histogram illustrating the distribution of TF binding targets detected by ChEC-seq. The x-axis represents the range of TFs binding targets and y-axis displays their frequency.

**(C)** Bar plots showing the distribution of TFIID and CR genes among binding targets of three representative TFs (Cyc8, YLL054C, and Mga1). TFIID promoters account for 87% of the genomic distribution, while CR promoters represent 13%. TFs with more than 87% TFIID genes as binding targets and p-value < 0.05 in hypergeometric test are considered enriched at TFIID promoters, and those with significantly more than 13% CR binding targets are enriched at CR promoters. TFs with similar distribution and no significant preference to either coactivator class are considered as unbiased.

**(D)** Boxplot illustrating the distribution of the number of TFs bound to CR (blue) and TFIID (orange) gene promoters.

**Figure S2. Comparison of TF binding measured by ChEC-seq and ChIP-exo**

**(A)** Protein coding gene promoters with detectable TF binding measured by ChIP-exo^30^ and where the TF DNA binding motif is known^35^. Y-axis shows data for the 68 TFs where binding has been mapped by both approaches. Green bars denote binding targets with DNA sequence motif; yellow bars denote binding targets without motif.

**(B)** Venn diagram showing overlap between ChEC-seq (blue) and ChIP-exo (orange) binding targets.

**(C)** Venn diagrams show overlap between ChEC-seq (blue) and ChIP-seq (orange) binding targets for Gal4 (following 120 min of galactose induction) and Gcn4 (following 60 min of SM induction).

**Figure S3. Properties of gene clusters based on the number of TF binding sites**

**(A)** Box plot displays the ChEC-seq DNA cleavage signal of TFs at promoters in each cluster. Average log2 values of ChEC signal around bound peaks from triplicates (−150 to +150 bps from center of peak) normalized to *Drosophila* spike-in DNA are plotted on the y-axis.

**(B)** Boxplot showing the expression levels of genes in each gene cluster (based on the number of TF binding sites from Fig 1B). The y-axis displays the range of expression values (log2 scale) in each cluster represented on the x-axis.

**(C)** Comparison of our TF binding clusters (Fig 1B) among previously published gene classes^30^ is shown.

**Figure S4. Activation domain of Gcn4 mediates its binding at Cluster-III promoters**

(A) Western blot analysis showing protein levels of Gcn4 (left), Gcn4-ad (middle) and Gcn4-dbd (right) in triplicates. The bottom blot shows signal for loading control Tfg2 (TFIIF subunit).

(B) A representative browser image is displayed to show the binding patterns of Gcn4 and its mutants. The binding patterns are shown for a portion of Chr-IV.

(C) Stacked bar plot displaying the total number of binding sites (y-axis) of Gcn4 and its mutants (x-axis). Binding sites with Gcn4 consensus motifs are depicted in green and without motif are denoted in yellow.

(D) The heatmap displays the binding patterns of Gcn4, Gcn4-ad, Gcn4-dbd, Med15, Swi3, Swp82, Bur6 and Mot1 to different clusters obtained in Figure 1. The binding patterns of Gcn4 and its mutants are shown in the absence and presence of SM stress for 60 min. The color bar on the right denotes genes that are present near chromosome domain (L-CID) boundaries (in blue) ^47^ across clusters. Each cluster is separated by a red line for easy visualization.

**Figure S5. Rapid TF depletion to identify expression targets**

**(A)** The Western blot analysis shows representative TF-degron protein levels following 30 min of IAA treatment for three replicates. The blot is probed with anti-V5 antibody to detect the degron tagged TF and the bottom blot shows the signal for the loading control Tfg2. (Complete data set available at <u>Mendeley link</u>).

**(B)** The left bar plot displays the number of genes affected at different time points of Reb1 depletion. The yellow bar represents genes located within 500 bps of Reb1 binding sites, the blue bar represents genes located within 10 kb distance, and the gray bar represents genes located further than 10 kb away. The right panel shows a representative browser shot of Reb1 binding sites in blue at the top of the panel and genes affected by its depletion at various time points shown in red. All protein coding genes in the region are shown at bottom.

**(C)** Boxplot shows the number of TFs that, when rapidly depleted, alter gene expression from individual TFIID and CR genes. A median of 10 TFs affects expression of CR genes, and 3 TFs affect TFIID genes. Welsh’s t-test results are shown.

**(D)** The boxplot shows the number of TFs that affect gene expression from genes in each gene cluster obtained by K-means clustering from Fig1B.

**Figure S6: The response towards TF depletion is comparable across gene clusters**

**(A)** Box plot displaying the median fold change in each cluster (based on **Fig 1B**). The y-axis shows range of log2 fold change values of genes that are significantly affected (**Methods**) upon depleting 126 TFs in each cluster obtained from **Fig 2B** (x-axis). The right panel shows transcription changes in response to factors that lead to a transcription decrease while the left panel shows the response to factor depletion that leads to a transcription increase.

**(B)** Same as **S6A** but range of log2 fold change values are shown for gene classes based on ChIP-exo TF binding data^30^.

**Figure S7. Merging ChIP-exo binding and expression data**

**(A)** Bar plot displays the fraction of functional TF binding sites among the total number of binding sites. The y-axis represents each TF and x-axis displays the number of binding targets determined by ChIP-exo^30^. The blue bar indicates the functional binding targets, while the yellow bar displays binding targets in gene regulatory regions where transcription is not altered by factor depletion.

**(B)** The bar plot displays the proportion of functional binding sites in relation to the total number of expression targets. The y-axis represents each transcription factor (TF) and x-axis displays the number of expression targets. The blue bars indicate the number of functional targets where the TF binding site (determined by ChIP-exo) lies in the regulatory region of the affected gene, and the pink bars represent expression targets without detectable TF binding measured by ChIP-exo.

**Figure S8. Functional targets are modestly enriched for TF binding sites but with many exceptions**

**(A)** Bar plot displays the number of expression targets for each analyzed TF. The blue bar represents the expression targets with corresponding TF motif in the gene regulatory region (−400 to +200 bps from TSS) and the green bar shows targets without the motif.

**(B)** Left panel illustrates the overlap between TF binding targets (measured by ChEC) and expression targets for 126 TFs. The right panel shows the overlap between binding targets (measured by ChIP-exo) and expression targets for 78 TFs.

**(C, D)** The scatter plot presents p-values of the hypergeometric test. For 73% of TFs measured (92/126), functional targets are significantly enriched in ChEC binding sites when compared to random chance (**C**). Similarly, for about 65% of TFs measured (51/78 TFs), functional targets are significantly enriched in ChIP-exo binding sites (**D**). TFs with significant p-value (<0.05) are represented below the red dotted line.

**(E)** Heatmaps showing percentage of functional binding targets in each coactivator class. Color bar goes from 0 to 60 percent (from yellow to red). A total of 116 TFs having at least one common binding and expression target are shown in two groups for easier visualization. Top row in each heatmap represents TFIID genes and bottom row is CR. Higher percent of CR binding targets are functional when compared to TFIID binding targets in the majority of cases (72%).

**Figure S9. Depletion of TFs under various stress conditions**

The differential expression of genes upon depletion of the indicated transcription factors is represented by volcano plots. Significantly affected genes are highlighted in red and the number of genes that are upregulated or downregulated in each case are denoted on the plot.

**(A)** The left plot shows the effect of Gcn4 depletion without SM stress, while the right plot displays the effect of Gcn4 depletion following 60 min of SM stress.

**(B)** The left plot shows the effect of Hsf1 depletion without heat shock, while the right plot displays the effect of Hsf1 depletion after 10 min heat shock at 37°C.

**(C)** The left plot shows the effect of Msn2 depletion without heat shock, while the right plot displays the effect of Msn2 depletion after 10 min heat shock at 37°C.

**(D)** The left plot displays genes affected by Gal4 depletion without galactose induction, while the right plot shows the effect of Gal4 depletion 2 hours after galactose induction.

**(E)** The bar plot displays the proportion of functional binding sites among the total number of expression targets. The y-axis represents each TF under specified conditions (**Fig S9A-D**) and x-axis displays the number of expression targets. The blue bars indicate the number of functional targets where the TF binding site lies in the regulatory region of the affected gene, and the pink bars represent expression targets without detectable TF binding measured by ChEC-seq.

**(F)** The bar plot displays the fraction of functional TF binding sites among the total number of binding sites. ChEC-seq of Gcn4 is performed in −/+ SM stress and Gal4 in −/+ galactose induction, non-heat shock binding profiles are used for comparisons in case of Hsf1 and Msn2. The blue bar indicates functional binding targets, while the yellow bar displays binding targets in gene regulatory regions where transcription of the downstream gene is not altered by factor depletion.

## References

1. Fong, A. P. & Tapscott, S. J. Skeletal muscle programming and re-programming. Curr Opin Genet Dev 23, 568–573 (2013).

2. Hahn, S. & Young, E. T. Transcriptional regulation in Saccharomyces cerevisiae: transcription factor regulation and function, mechanisms of initiation, and roles of activators and coactivators. Genetics 189, 705–736 (2011).

3. Lambert, S. A. et al. The Human Transcription Factors. Cell 172, 650–665 (2018).

4. Lee, T. I. & Young, R. A. Transcriptional regulation and its misregulation in disease. Cell 152, 1237–1251 (2013).

5. Singh, H., Khan, A. A. & Dinner, A. R. Gene regulatory networks in the immune system. Trends Immunol 35, 211–218 (2014).

6. Takahashi, K. & Yamanaka, S. A decade of transcription factor-mediated reprogramming to pluripotency. Nat Rev Mol Cell Biol 17, 183–193 (2016).

7. Soto, L. F. et al. Compendium of human transcription factor effector domains. Mol Cell 82, 514–526 (2022).

8. Lu, F. & Lionnet, T. Transcription Factor Dynamics. Cold Spring Harb Perspect Biol 13, (2021).

9. Kribelbauer, J. F., Rastogi, C., Bussemaker, H. J., Mann, R. S. & Zuckerman, M. B. Low-Affinity Binding Sites and the Transcription Factor Specificity Paradox in Eukaryotes. Annual Review of Cell and Developmental Biology Annu. Rev. Cell Dev. Biol 35, 357–79 (2019).

10. Smith, N. C. & Matthews, J. M. Mechanisms of DNA-binding specificity and functional gene regulation by transcription factors. Curr Opin Struct Biol 38, 68–74 (2016).

11. Slattery, M. et al. Absence of a simple code: how transcription factors read the genome. Trends Biochem Sci 39, 381–399 (2014).

12. Brodsky, S., et al. Intrinsically Disordered Regions Direct Transcription Factor In Vivo Binding Specificity. Mol Cell 79, 459–471.e4 (2020).

13. Divya Krishna Kumar, A., et al. Complementary strategies for directing in vivo transcription factor binding through DNA binding domains and intrinsically disordered regions. (2023) doi:10.1016/j.molcel.2023.04.002.

14. Jolma, A. et al. DNA-dependent formation of transcription factor pairs alters their binding specificity. Nature 527, 384–388 (2015).

15. Slattery, M. et al. Cofactor Binding Evokes Latent Differences in DNA Binding Specificity between Hox Proteins. Cell 147, 1270–1282 (2011).

16. Alerasool, N., Leng, H., Lin, Z.-Y., Gingras, A.-C. & Taipale, M. Identification and functional characterization of transcriptional activators in human cells. Mol Cell (2022) doi:10.1016/j.molcel.2021.12.008 PMID - 35016035.

17. Erijman, A. et al. A High-Throughput Screen for Transcription Activation Domains Reveals Their Sequence Features and Permits Prediction by Deep Learning. Mol Cell 78, 890–902.e6 (2020).

18. Sanborn, A. L. et al. Simple biochemical features underlie transcriptional activation domain diversity and dynamic, fuzzy binding to Mediator. Elife 10, e68068 (2021).

19. Sahu, B. et al. Sequence determinants of human gene regulatory elements. Nat Genet 54, 283–294 (2022).

20. Panne, D. The enhanceosome. Curr Opin Struct Biol 18, 236–242 (2008).

21. Kulkarni, M. M. & Arnosti, D. N. Information display by transcriptional enhancers. Development 130, 6569–6575 (2003).

22. King, D. M. et al. Synthetic and genomic regulatory elements reveal aspects of cis-regulatory grammar in Mouse Embryonic Stem Cells. Elife 9, e41279 (2020).

23. de Boer, C. G. et al. Deciphering eukaryotic gene-regulatory logic with 100 million random promoters. Nat Biotechnol 38, 56–65 (2020).

24. Donczew, R. & Hahn, S. Bet family members bdf1/2 modulate global transcription initiation and elongation in saccharomyces cerevisiae. Elife 10, (2021).

25. Warfield, L., Donczew, R., Mahendrawada, L. & Hahn, S. Yeast Mediator facilitates transcription initiation at most promoters via a Tail-independent mechanism. Mol Cell 82, 4033–4048.e7 (2022).

26. Donczew, R., Warfield, L., Pacheco, D., Erijman, A. & Hahn, S. Two roles for the yeast transcription coactivator SAGA and a set of genes redundantly regulated by TFIID and SAGA. Elife 9, (2020).

27. Schofield, J. A. & Hahn, S. Broad compatibility between yeast UAS elements and core promoters and identification of promoter elements that determine cofactor specificity. CellReports 42, 112387 (2023).

28. Chua, G., et al. Identifying transcription factor functions and targets by phenotypic activation. (2006).

29. Hu, Z., Killion, P. J. & Iyer, V. R. Genetic reconstruction of a functional transcriptional regulatory network. Nature Genetics 2007 39:5 39, 683–687 (2007).

30. Rossi, M. J. et al. A high-resolution protein architecture of the budding yeast genome. Nature 592, 309–314 (2021).

31. Sopko, R. et al. Mapping Pathways and Phenotypes by Systematic Gene Overexpression. Mol Cell 21, 319–330 (2006).

32. Zentner, G. E., Kasinathan, S., Xin, B., Rohs, R. & Henikoff, S. ChEC-seq kinetics discriminates transcription factor binding sites by DNA sequence and shape in vivo. (2015) doi:10.1038/ncomms9733.

33. Donczew, R., Lalou, A., Devys, D., Tora, L. & Hahn, S. An improved ChEC-seq method accurately maps the genome-wide binding of transcription coactivators and sequence-specific transcription factors. bioRxiv 2021.02.12.430999 (2021) doi:10.1101/2021.02.12.430999.

34. Huisinga, K. L. & Pugh, B. F. A Genome-Wide Housekeeping Role for TFIID and a Highly Regulated Stress-Related Role for SAGA in Saccharomyces cerevisiae. Molecular Cell vol. 13 (2004).

35. Castro-Mondragon, J. A. et al. JASPAR 2022: the 9th release of the open-access database of transcription factor binding profiles. Nucleic Acids Res 50, D165–D173 (2022).

36. Grant, C. E., Bailey, T. L. & Noble, W. S. Sequence analysis FIMO: scanning for occurrences of a given motif. Bioinformatics 27, 1017–1018 (2011).

37. Chereji, R. V., Ramachandran, S., Bryson, T. D. & Henikoff, S. Precise genome-wide mapping of single nucleosomes and linkers in vivo. Genome Biol 19, 1–20 (2018).

38. Nguyen, V. Q. et al. Spatiotemporal coordination of transcription preinitiation complex assembly in live cells. Mol Cell 81, 3560–3575.e6 (2021).

39. Kim, J. M. et al. Single-molecule imaging of chromatin remodelers reveals role of ATPase in promoting fast kinetics of target search and dissociation from chromatin. Elife 10, (2021).

40. Mehta, G. D. et al. Single-Molecule Analysis Reveals Linked Cycles of RSC Chromatin Remodeling and Ace1p Transcription Factor Binding in Yeast In Brief Article Single-Molecule Analysis Reveals Linked Cycles of RSC Chromatin Remodeling and Ace1p Transcription Factor Binding in Yeast. doi:10.1016/j.molcel.2018.09.009.

41. Bruzzone, M. J., Grünberg, S., Kubik, S., Zentner, G. E. & Shore, D. Distinct patterns of histone acetyltransferase and mediator deployment at yeast protein-coding genes. Genes Dev 32, 1252–1265 (2018).

42. Ellenberger, T. E., Brandl, C. J., Struhl, K. & Harrison, S. C. The GCN4 basic region leucine zipper binds DNA as a dimer of uninterrupted α Helices: Crystal structure of the protein-DNA complex. Cell 71, 1223–1237 (1992).

43. Kim, J. & Struhl, K. Determinants of half-site spacing preferences that distinguish AP-1 and ATF/CREB bZIP domains. Nucleic Acids Res 23, 2531–2537 (1995).

44. Seong, K. M. et al. A new method for the construction of a mutant library with a predictable occurrence rate using Poisson distribution. J Microbiol Methods 69, 442–450 (2007).

45. Brzovic, P. S. et al. The acidic transcription activator Gcn4 binds the mediator subunit Gal11/Med15 using a simple protein interface forming a fuzzy complex. Mol Cell 44, 942– 953 (2011).

46. Drysdale, C. M. et al. The transcriptional activator GCN4 contains multiple activation domains that are critically dependent on hydrophobic amino acids. Mol Cell Biol 15, 1220–1233 (1995).

47. Swygert, S. G. et al. Condensin-Dependent Chromatin Compaction Represses Transcription Globally during Quiescence. Mol Cell 73, 533–546.e4 (2019).

48. Chereji, R. V. et al. Mediator binds to boundaries of chromosomal interaction domains and to proteins involved in DNA looping, RNA metabolism, chromatin remodeling, and actin assembly. Nucleic Acids Res 45, 8806 (2017).

49. Hsieh, T. H. S. et al. Mapping Nucleosome Resolution Chromosome Folding in Yeast by Micro-C. Cell 162, 108–119 (2015).

50. Nishimura, K., Fukagawa, T., Takisawa, H., Kakimoto, T. & Kanemaki, M. An auxin-based degron system for the rapid depletion of proteins in nonplant cells. Nature Methods 2009 6:12 6, 917–922 (2009).

51. Chan, L. Y., Mugler, C. F., Heinrich, S., Vallotton, P. & Weis, K. Non-invasive measurement of mRNA decay reveals translation initiation as the major determinant of mRNA stability. Elife 7, (2018).

52. Hartley, P. D. & Madhani, H. D. Mechanisms that Specify Promoter Nucleosome Location and Identity. Cell 137, 445–458 (2009).

53. Yan, C., Chen, H. & Bai, L. Systematic Study of Nucleosome-Displacing Factors in Budding Yeast. Mol Cell 71, 294–305.e4 (2018).

54. Fourel, G. et al. An activation-independent role of transcription factors in insulator function. EMBO Rep 2, 124 (2001).

55. Cherry, J. M. et al. Saccharomyces Genome Database: the genomics resource of budding yeast. Nucleic Acids Res 40, D700 (2012).

56. Love, M. I., Huber, W. & Anders, S. Moderated estimation of fold change and dispersion for RNA-seq data with DESeq2. Genome Biol 15, 1–21 (2014).

57. Thomas, P. D. et al. PANTHER: Making genome-scale phylogenetics accessible to all. Protein Sci 31, 8–22 (2022).

58. Chen, H. & Pugh, B. F. What do Transcription Factors Interact With? J Mol Biol 433, (2021).

59. Baker Brachmann, C., et al. Designer deletion strains derived fromSaccharomyces cerevisiae S288C: A useful set of strains and plasmids for PCR-mediated gene disruption and other applications. Yeast 14, 115–132 (1998).

60. Grünberg, S., Henikoff, S., Hahn, S. & Zentner, G. E. Mediator binding to UAS s is broadly uncoupled from transcription and cooperative with TFIID recruitment to promoters . EMBO J 35, 2435–2446 (2016).

61. Warfield, L. et al. Transcription of Nearly All Yeast RNA Polymerase II-Transcribed Genes Is Dependent on Transcription Factor TFIID. Mol Cell 68, 118–129.e5 (2017).

62. Bonnet, J. et al. The SAGA coactivator complex acts on the whole transcribed genome and is required for RNA polymerase II transcription. Genes Dev 28, 1999–2012 (2014).

63. Duffy, E. E. et al. Tracking Distinct RNA Populations Using Efficient and Reversible Covalent Chemistry. Mol Cell 59, 858–866 (2015).

64. Duffy, E. E. & Simon, M. D. Enriching s ^4^ U-RNA Using Methane Thiosulfonate (MTS) Chemistry. Curr Protoc Chem Biol 8, 234–250 (2016).

65. Langmead, B. & Salzberg, S. L. Fast gapped-read alignment with Bowtie 2. Nat Methods 9, 357–359 (2012).

66. Heinz, S. et al. Simple Combinations of Lineage-Determining Transcription Factors Prime cis-Regulatory Elements Required for Macrophage and B Cell Identities. Mol Cell 38, 576– 589 (2010).

67. Park, D., Morris, A. R., Battenhouse, A. & Iyer, V. R. Simultaneous mapping of transcript ends at single-nucleotide resolution and identification of widespread promoter-associated non-coding RNA governed by TATA elements. Nucleic Acids Res 42, 3736–3749 (2014).

68. Anders, S., Pyl, P. T. & Huber, W. HTSeq—a Python framework to work with high-throughput sequencing data. Bioinformatics 31, 166–169 (2015).

69. Love, M. I., Huber, W. & Anders, S. Moderated estimation of fold change and dispersion for RNA-seq data with DESeq2. Genome Biol 15, 550 (2014).

