## supplemental figures 1-9 for "Surprising connections between DNA binding and function for the near-complete set of yeast transcription factors"

S1

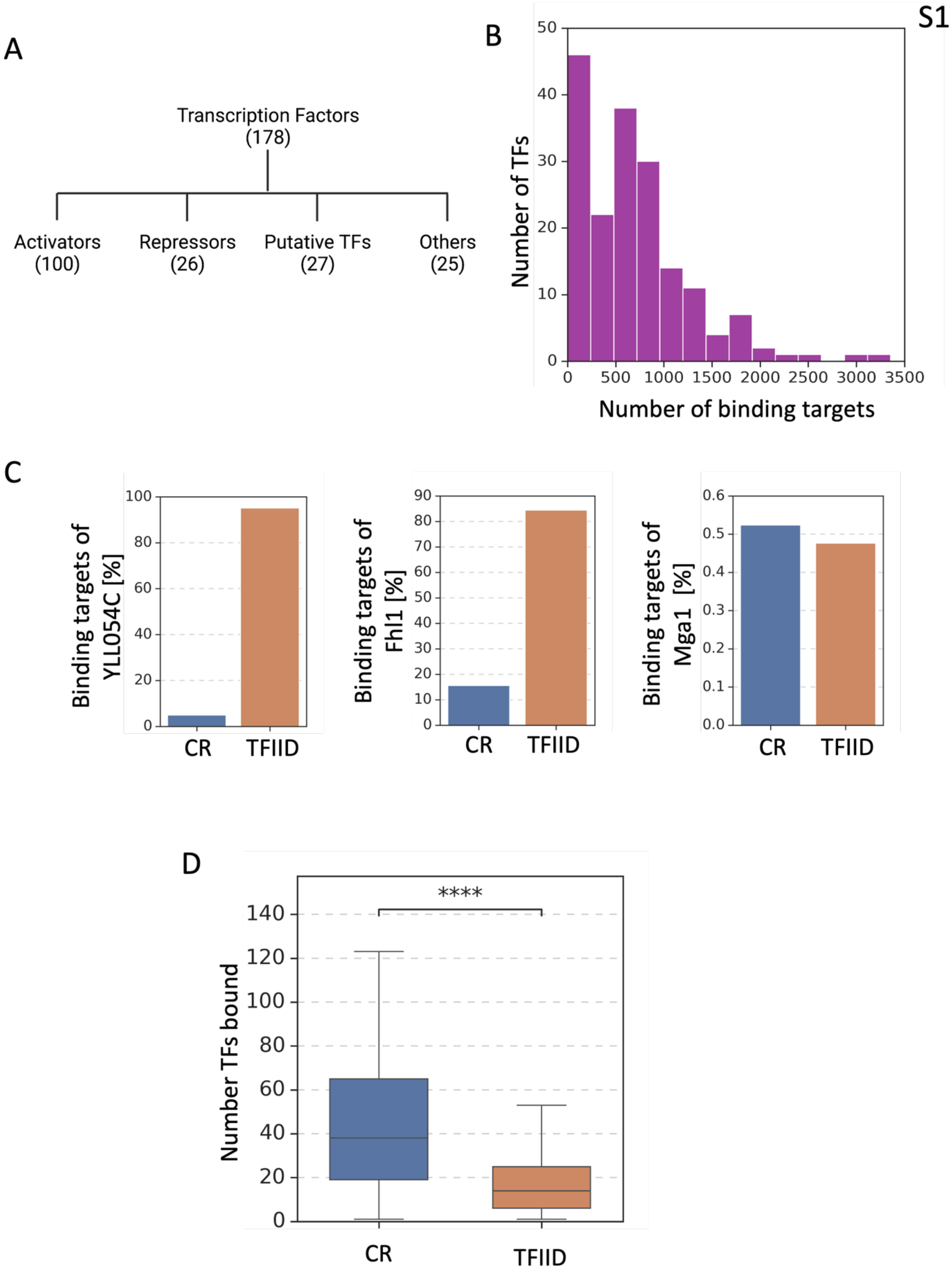
A
Transcription Factors
(178)
Activators
(100)
Repressors
(26)
Putative TFs
(27)
Others
(25)
B
Number of TFs
Number of binding targets
0
500
1000
1500
2000
2500
3000
3500
10
20
30
40
50
C
Binding targets of YLL054C [%]
Binding targets of Fhl1 [%]
Binding targets of Mga1 [%]
CR
TFIID
D
Number TFs bound
****
CR
TFIID

**Figure S1. ChEC-seq mapping of yeast transcription factors (TFs)**
**(A)** Schematic showing the functional categories of 178 TFs with DNA binding successfully analyzed by ChEC-seq.
**(B)** Histogram illustrating the distribution of TF binding targets detected by ChEC-seq. The x-axis represents the range of TFs binding targets and y-axis displays their frequency.
**(C)** Bar plots showing the distribution of TFIID and CR genes among binding targets of three representative TFs (Cyc8, YLL054C, and Mga1). TFIID promoters account for 87% of the genomic distribution, while CR promoters represent 13%. TFs with more than 87% TFIID genes as binding targets and p-value $< 0.05$ in hypergeometric test are considered enriched at TFIID promoters, and those with significantly more than 13% CR binding targets are enriched at CR promoters. TFs with similar distribution and no significant preference to either coactivator class are considered as unbiased.
**(D)** Boxplot illustrating the distribution of the number of TFs bound to CR (blue) and TFIID (orange) gene promoters.

S2

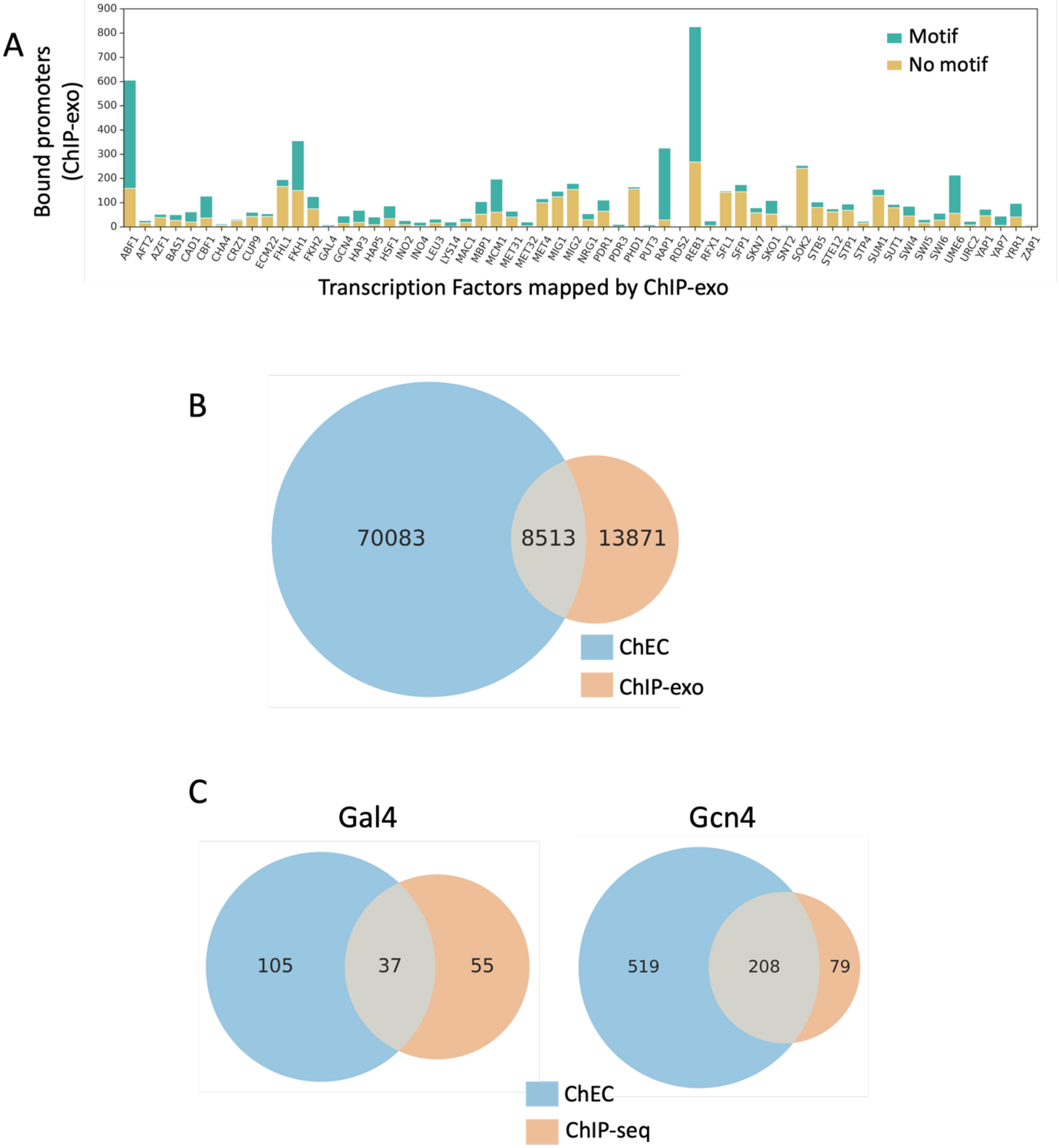

A
Bound promoters (ChIP-exo)
900
800
700
600
500
400
300
200
100
0
Motif
No motif
ABF1 AFT2 AZF1 BAS1 CAD1 CBF1 CHA4 CRZ1 CUP9 ECM22 FHL1 FKH1 FKH2 GAL4 GCN4 HAP3 HAP5 HSF1 INO2 INO4 LEU3 LYS14 MAC1 MBP1 MCM1 MET31 MET32 MET4 MIG1 MIG2 NRG1 PDR1 PDR3 PHD1 PUT3 RAP1 RDS2 REB1 RFX1 SFL1 SFP1 SKN7 SKO1 SNT2 SOK2 STB5 STE12 STP1 STP4 SUM1 SUT1 SWI4 SWI5 SWI6 UME6 URC2 YAP1 YAP7 YRR1 ZAP1
Transcription Factors mapped by ChIP-exo
B
70083
8513
13871
ChEC
ChIP-exo
C
Gal4
105
37
55
Gcn4
519
208
79
ChEC
ChIP-seq

**Figure S2. Comparison of TF binding measured by ChEC-seq and ChIP-exo**

**(A)** Protein coding gene promoters with detectable TF binding measured by ChIP-exo[30] and where the TF DNA binding motif is known[35]. Y-axis shows data for the 68 TFs where binding has been mapped by both approaches. Green bars denote binding targets with DNA sequence motif; yellow bars denote binding targets without motif.

**(B)** Venn diagram showing overlap between ChEC-seq (blue) and ChIP-exo (orange) binding targets.

**(C)** Venn diagrams show overlap between ChEC-seq (blue) and ChIP-seq (orange) binding targets for Gal4 (following 120 min of galactose induction) and Gcn4 (following 60 min of SM induction).

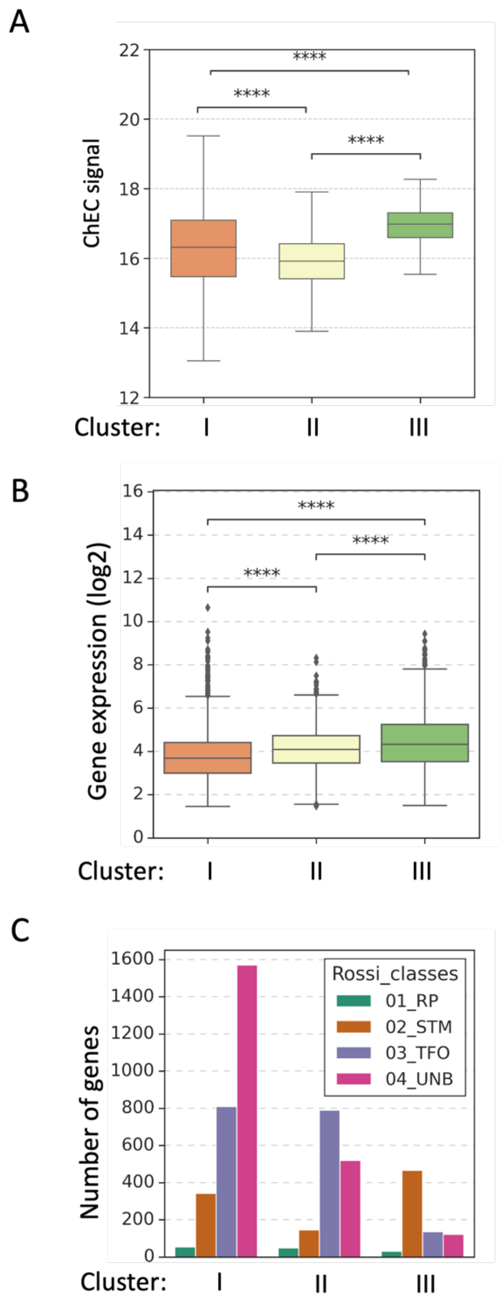


**Figure S3. Properties of gene clusters based on the number of TF binding sites**

**(A)** Box plot displays the ChEC-seq DNA cleavage signal of TFs at promoters in each cluster. Average log2 values of ChEC signal around bound peaks from triplicates (-150 to +150 bps from center of peak) normalized to *Drosophila* spike-in DNA are plotted on the y-axis.

**(B)** Boxplot showing the expression levels of genes in each gene cluster (based on the number of TF binding sites from Fig 1B). The y-axis displays the range of expression values (log2 scale) in each cluster represented on the x-axis.

**(C)** Comparison of our TF binding clusters (Fig 1B) among previously published gene classes[30] is shown.

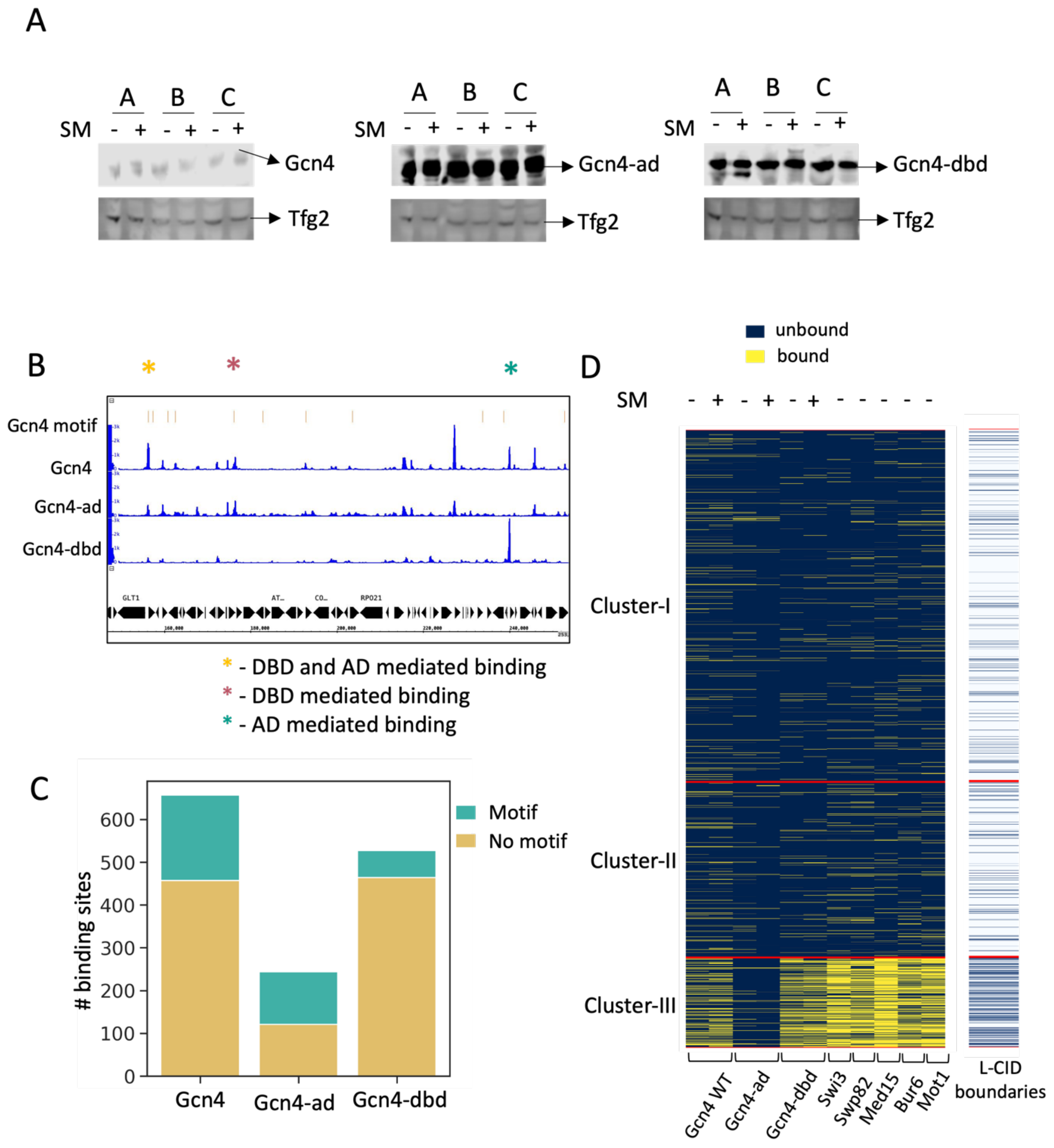
A
A B C
SM - + - + - +
Gcn4
Tfg2
Gcn4-ad
Gcn4-dbd
B
Gcn4 motif
Gcn4
Gcn4-ad
Gcn4-dbd
GLT1
RPO21
* - DBD and AD mediated binding
* - DBD mediated binding
* - AD mediated binding
C
### binding sites
0
100
200
300
400
500
600
Motif
No motif
Gcn4
Gcn4-ad
Gcn4-dbd
D
unbound
bound
SM - + - + - + - - - - -
Cluster-I
Cluster-II
Cluster-III
Gcn4 WT
Gcn4-ad
Gcn4-dbd
Swi3
Swp82
Med15
Bur6
Mot1
L-CID boundaries

**Figure S4. Activation domain of Gcn4 mediates its binding at Cluster-III promoters**

(**A**) Western blot analysis showing protein levels of Gcn4 (left), Gcn4-ad (middle) and Gcn4-dbd (right) in triplicates. The bottom blot shows signal for loading control Tfg2 (TFIIF subunit).

(**B**) A representative browser image is displayed to show the binding patterns of Gcn4 and its mutants. The binding patterns are shown for a portion of Chr-IV.

(**C**) Stacked bar plot displaying the total number of binding sites (y-axis) of Gcn4 and its mutants (x-axis). Binding sites with Gcn4 consensus motifs are depicted in green and without motif are denoted in yellow.

(**D**) The heatmap displays the binding patterns of Gcn4, Gcn4-ad, Gcn4-dbd, Med15, Swi3, Swp82, Bur6 and Mot1 to different clusters obtained in Figure 1. The binding patterns of Gcn4 and its mutants are shown in the absence and presence of SM stress for 60 min. The color bar on the right denotes genes that are present near chromosome domain (L-CID) boundaries (in blue) [47] across clusters. Each cluster is separated by a red line for easy visualization.

S5

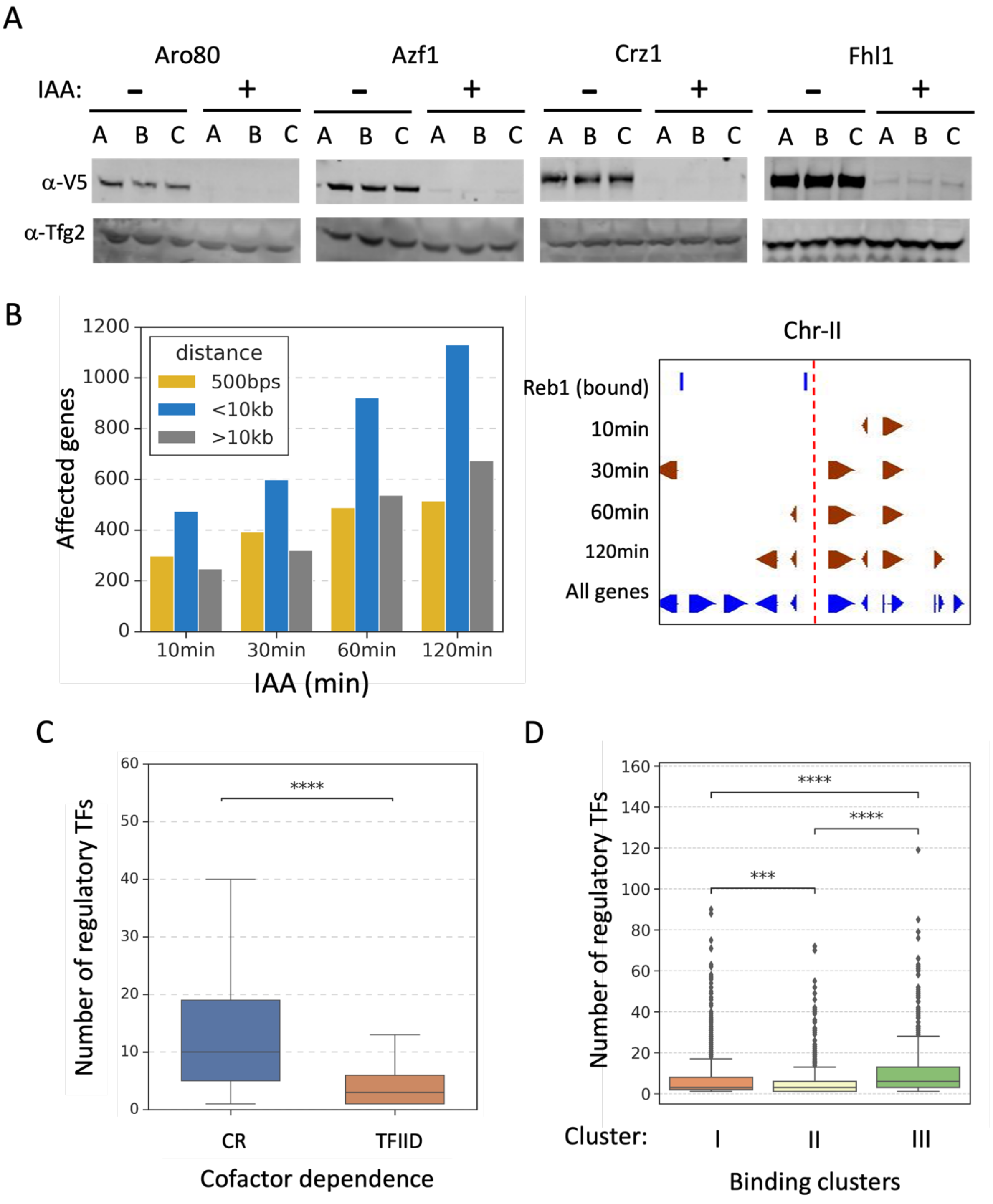
A
Aro80
Azf1
Crz1
Fhl1
IAA:
− +
A B C A B C
α-V5
α-Tfg2
B
Affected genes
distance
500bps
<10kb
>10kb
0
200
400
600
800
1000
1200
10min
30min
60min
120min
IAA (min)
Chr-II
Reb1 (bound)
10min
30min
60min
120min
All genes
C
Number of regulatory TFs
****
CR
TFIID
Cofactor dependence
D
Number of regulatory TFs
****
***
Cluster:
I
II
III
Binding clusters

**Figure S5. Rapid TF depletion to identify expression targets**

**(A)** The Western blot analysis shows representative TF-degron protein levels following 30 min of IAA treatment for three replicates. The blot is probed with anti-V5 antibody to detect the degron tagged TF and the bottom blot shows the signal for the loading control Tfg2. (Complete data set available at Mendeley link).

**(B)** The left bar plot displays the number of genes affected at different time points of Reb1 depletion. The yellow bar represents genes located within 500 bps of Reb1 binding sites, the blue bar represents genes located within 10 kb distance, and the gray bar represents genes located further than 10 kb away. The right panel shows a representative browser shot of Reb1 binding sites in blue at the top of the panel and genes affected by its depletion at various time points shown in red. All protein coding genes in the region are shown at bottom.

**(C)** Boxplot shows the number of TFs that, when rapidly depleted, alter gene expression from individual TFIID and CR genes. A median of 10 TFs affects expression of CR genes, and 3 TFs affect TFIID genes. Welsh's t-test results are shown.

**(D)** The boxplot shows the number of TFs that affect gene expression from genes in each gene cluster obtained by K-means clustering from Fig1B.

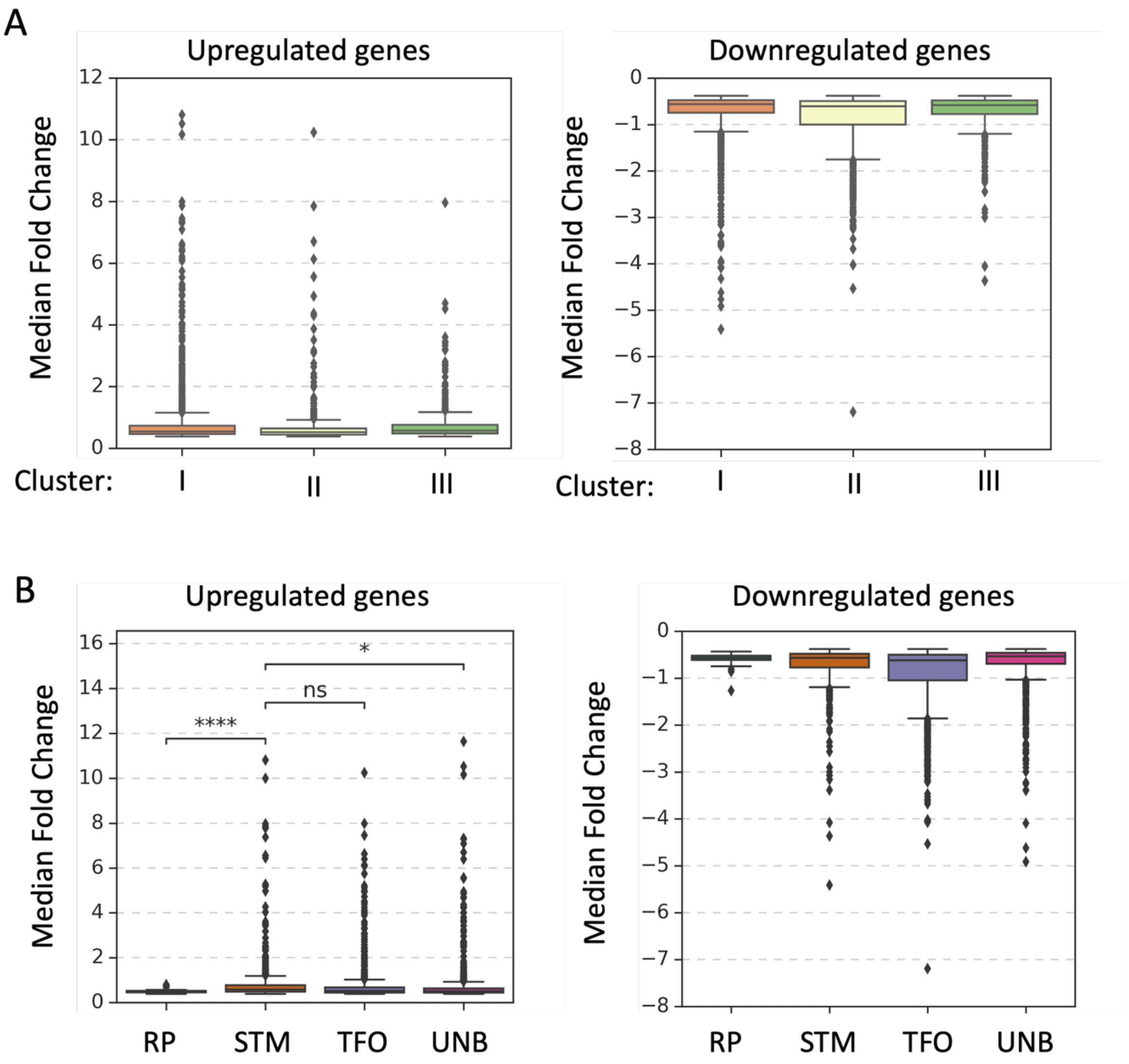


**Figure S6: The response towards TF depletion is comparable across gene clusters**

**(A)** Box plot displaying the median fold change in each cluster (based on **Fig 1B**). The y-axis shows range of log2 fold change values of genes that are significantly affected (**Methods**) upon depleting 126 TFs in each cluster obtained from **Fig 2B** (x-axis). The right panel shows transcription changes in response to factors that lead to a transcription decrease while the left panel shows the response to factor depletion that leads to a transcription increase.

**(B)** Same as **S6A** but range of log2 fold change values are shown for gene classes based on ChIP-exo TF binding data[30].

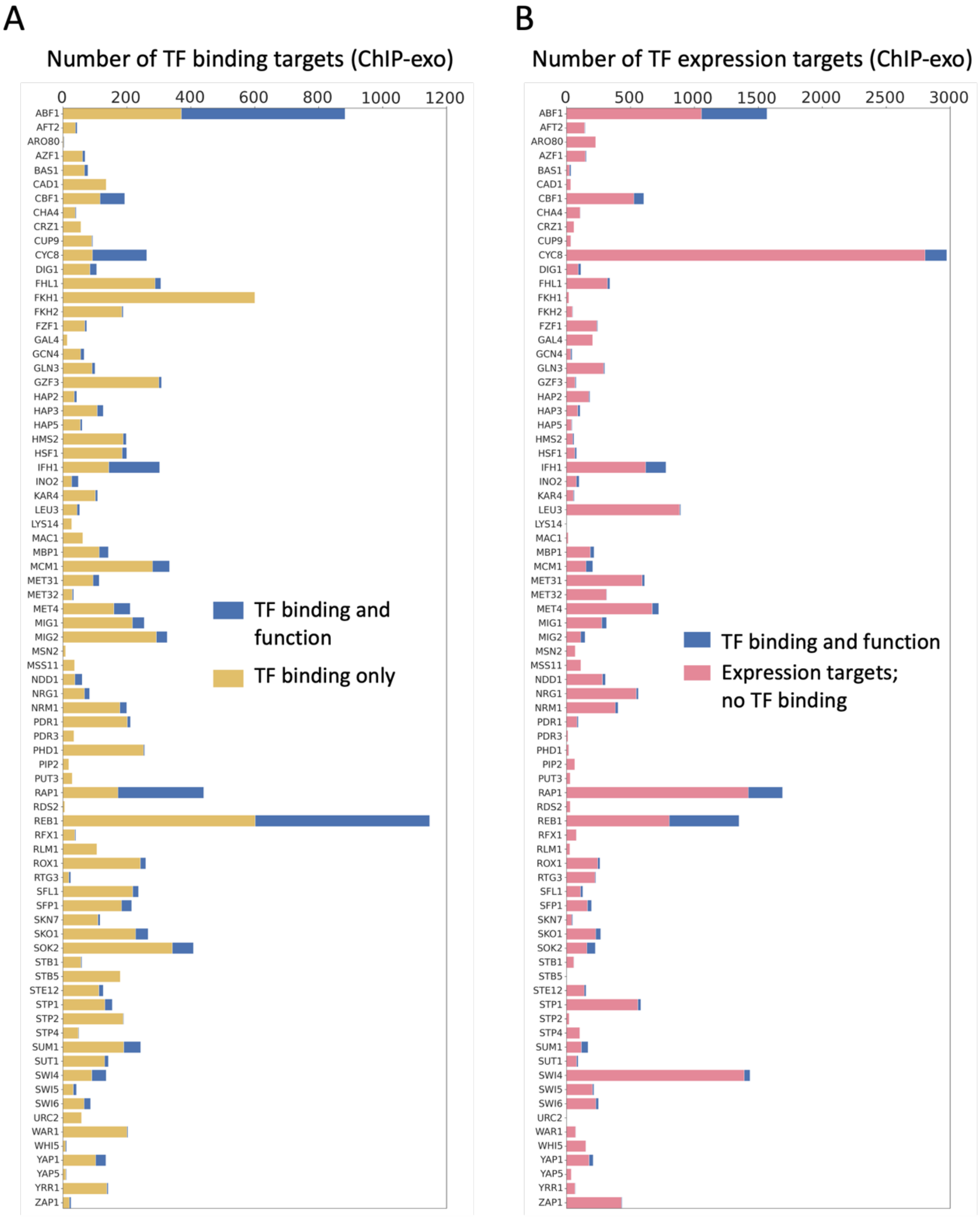
A
Number of TF binding targets (ChIP-exo)
0
200
400
600
800
1000
1200
TF binding and function
TF binding only
B
Number of TF expression targets (ChIP-exo)
0
500
1000
1500
2000
2500
3000
TF binding and function
Expression targets; no TF binding
ABF1
AFT2
ARO80
AZF1
BAS1
CAD1
CBF1
CHA4
CRZ1
CUP9
CYC8
DIG1
FHL1
FKH1
FKH2
FZF1
GAL4
GCN4
GLN3
GZF3
HAP2
HAP3
HAP5
HMS2
HSF1
IFH1
INO2
KAR4
LEU3
LYS14
MAC1
MBP1
MCM1
MET31
MET32
MET4
MIG1
MIG2
MSN2
MSS11
NDD1
NRG1
NRM1
PDR1
PDR3
PHD1
PIP2
PUT3
RAP1
RDS2
REB1
RFX1
RLM1
ROX1
RTG3
SFL1
SFP1
SKN7
SKO1
SOK2
STB1
STB5
STE12
STP1
STP2
STP4
SUM1
SUT1
SWI4
SWI5
SWI6
URC2
WAR1
WHI5
YAP1
YAP5
YRR1
ZAP1

**Figure S7. Merging ChIP-exo binding and expression data**

**(A)** Bar plot displays the fraction of functional TF binding sites among the total number of binding sites. The y-axis represents each TF and x-axis displays the number of binding targets determined by ChIP-exo[30]. The blue bar indicates the functional binding targets, while the yellow bar displays binding targets in gene regulatory regions where transcription is not altered by factor depletion.

**(B)** The bar plot displays the proportion of functional binding sites in relation to the total number of expression targets. The y-axis represents each transcription factor (TF) and x-axis displays the number of expression targets. The blue bars indicate the number of functional targets where the TF binding site (determined by ChIP-exo) lies in the regulatory region of the affected gene, and the pink bars represent expression targets without detectable TF binding measured by ChIP-exo.

S8

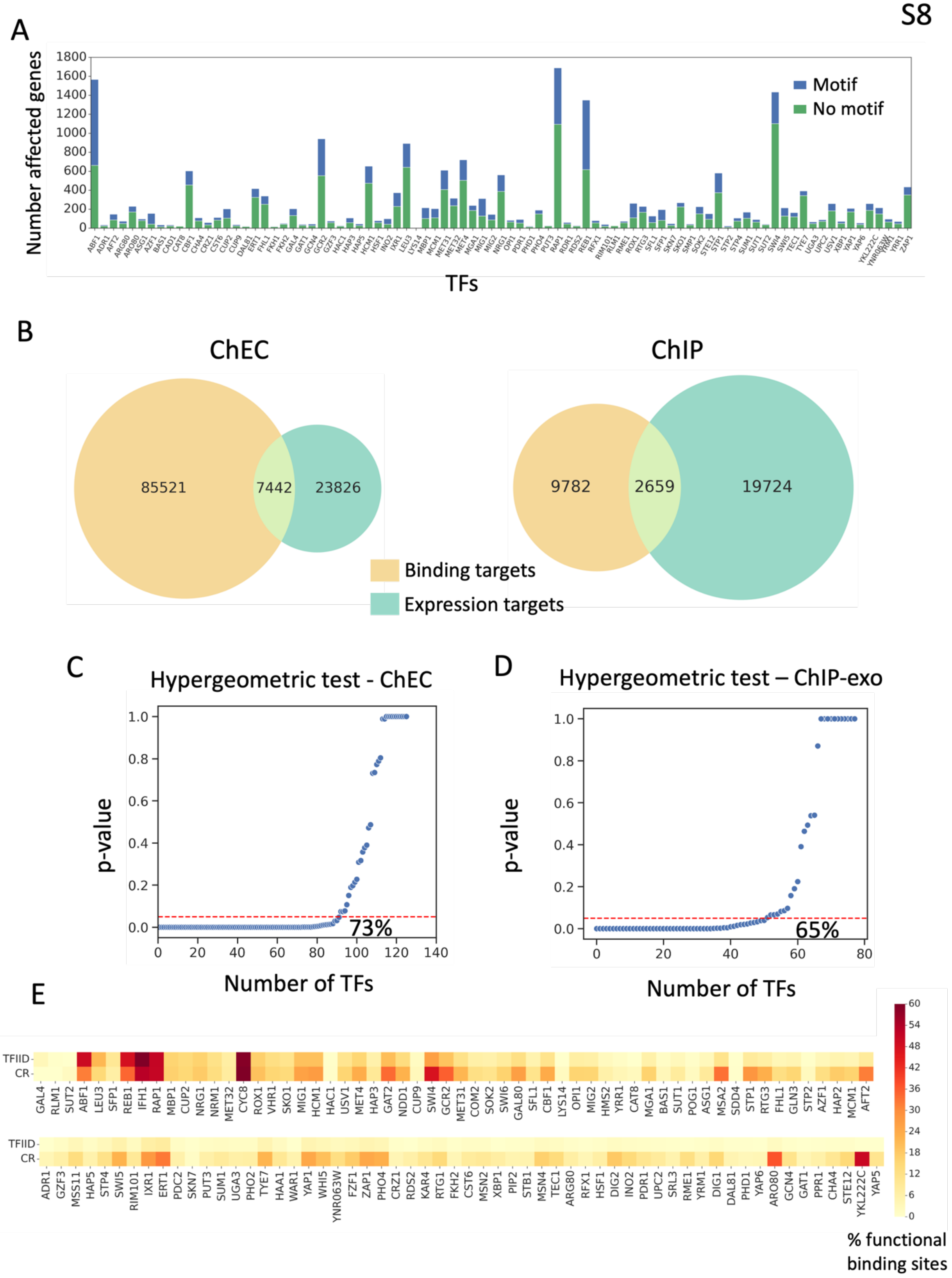
A
Number affected genes
TFs
Motif
No motif
B
ChEC
ChIP
85521
7442
23826
9782
2659
19724
Binding targets
Expression targets
C
Hypergeometric test - ChEC
p-value
Number of TFs
73%
D
Hypergeometric test – ChIP-exo
p-value
Number of TFs
65%
E
TFIID
CR
% functional binding sites

**Figure S8. Functional targets are modestly enriched for TF binding sites but with many exceptions**
**(A)** Bar plot displays the number of expression targets for each analyzed TF. The blue bar represents the expression targets with corresponding TF motif in the gene regulatory region (-400 to +200 bps from TSS) and the green bar shows targets without the motif.
**(B)** Left panel illustrates the overlap between TF binding targets (measured by ChEC) and expression targets for 126 TFs. The right panel shows the overlap between binding targets (measured by ChIP-exo) and expression targets for 78 TFs.
**(C, D)** The scatter plot presents p-values of the hypergeometric test. For 73% of TFs measured (92/126), functional targets are significantly enriched in ChEC binding sites when compared to random chance (**C**). Similarly, for about 65% of TFs measured (51/78 TFs), functional targets are significantly enriched in ChIP-exo binding sites (**D**). TFs with significant p-value ($<0.05$) are represented below the red dotted line.
**(E)** Heatmaps showing percentage of functional binding targets in each coactivator class. Color bar goes from 0 to 60 percent (from yellow to red). A total of 116 TFs having at least one common binding and expression target are shown in two groups for easier visualization. Top row in each heatmap represents TFIID genes and bottom row is CR. Higher percent of CR binding targets are functional when compared to TFIID binding targets in the majority of cases (72%).

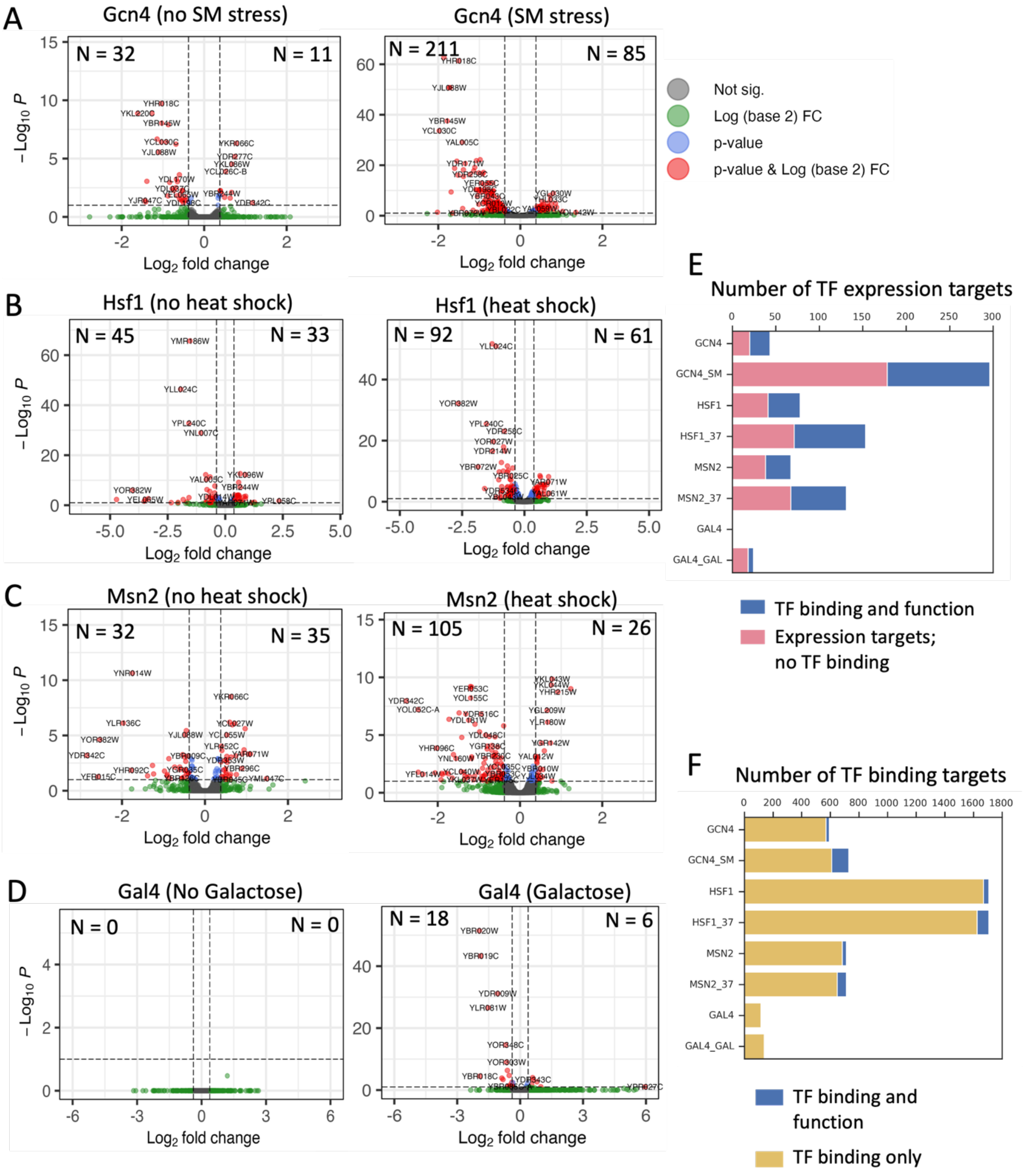


**Figure S9. Depletion of TFs under various stress conditions**

The differential expression of genes upon depletion of the indicated transcription factors is represented by volcano plots. Significantly affected genes are highlighted in red and the number of genes that are upregulated or downregulated in each case are denoted on the plot.

**(A)** The left plot shows the effect of Gcn4 depletion without SM stress, while the right plot displays the effect of Gcn4 depletion following 60 min of SM stress.
**(B)** The left plot shows the effect of Hsf1 depletion without heat shock, while the right plot displays the effect of Hsf1 depletion after 10 min heat shock at 37°C.
**(C)** The left plot shows the effect of Msn2 depletion without heat shock, while the right plot displays the effect of Msn2 depletion after 10 min heat shock at 37°C.
**(D)** The left plot displays genes affected by Gal4 depletion without galactose induction, while the right plot shows the effect of Gal4 depletion 2 hours after galactose induction.
**(E)** The bar plot displays the proportion of functional binding sites among the total number of expression targets. The y-axis represents each TF under specified conditions (**Fig S9A-D**) and x-axis displays the number of expression targets. The blue bars indicate the number of functional targets where the TF binding site lies in the regulatory region of the affected gene, and the pink bars represent expression targets without detectable TF binding measured by ChEC-seq.
**(F)** The bar plot displays the fraction of functional TF binding sites among the total number of binding sites. ChEC-seq of Gcn4 is performed in -/+ SM stress and Gal4 in -/+ galactose induction, non-heat shock binding profiles are used for comparisons in case of Hsf1 and Msn2. The blue bar indicates functional binding targets, while the yellow bar displays binding targets in gene regulatory regions where transcription of the downstream gene is not altered by factor depletion.
